# Inflammatory suppression and cancer-associated bypass of the b^0,+^ selenium-utilization program in the human intestine

**DOI:** 10.64898/2026.07.23.738870

**Authors:** Xiaobai He, Kundu Zhong, Yiyi Pu, Hui Hu, Yaoqiang Du, Xiaopan Chen, Linjie Chen

## Abstract

The b^0,+^ transporter (*SLC3A1*/*SLC7A9*) imports cystine and selenocystine at the enterocyte apical membrane and defines a selenium-utilization program in the healthy human intestine (He et al., *bioRxiv* 2026); how this program is altered in disease is unknown. Using a multi-cohort single-cell framework with donor-level statistics, we find that inflammation and cancer remodel a single epithelial redox-currency axis in opposite directions. In adult Crohn’s disease, b^0,+^ co-expression in small-intestinal enterocytes was suppressed (median 4.9% vs 34% positive; *P* = 0.008) but preserved in paediatric IBD; this loss reflected replacement of b^0,+^-high mature enterocytes by a dedifferentiated state rather than transcriptional disruption, with the *SLC7A9*-*SELENOP* coupling intact. Downstream, *GPX4* alone was selectively suppressed while the selenocysteine-incorporation machinery was preserved, priming the epithelium for ferroptosis in a severity-graded manner. Inflammatory cytokines (IFNγ, TNF) suppressed the selenium pole in small-intestinal enteroids, and a network knockout indicated that b^0,+^ supports the selenoproteome through substrate supply rather than transcriptional control; an independent patient proteome and a Caco-2 polarization model corroborated the axis. Colorectal-cancer colonocytes showed the opposing pole, near-absent b^0,+^ with induction of the xCT/thiol program (*P* = 5.6 × 10^−5^) and a ferroptosis-resistant configuration. Thus b^0,+^ marks the supply node of a dual-pole selenium-thiol axis: inflammation collapses the selenium pole toward a ferroptosis-prone dedifferentiated state, whereas cancer bypasses it toward an xCT antioxidant pole. The baseline axis state is an exploratory correlate of anti-TNF response and a candidate point of disease-associated vulnerability.

## Introduction

Selenium is an essential micronutrient that acts largely through its incorporation, as the amino acid selenocysteine, into roughly 25 selenoproteins, several of which, notably the glutathione peroxidases, defend cells against oxidative and lipid-peroxidative injury (Rayman, *Lancet* 2012). In the intestine, selenium and antioxidant selenoproteins shape both epithelial and immune homeostasis, and their supply is frequently disturbed in disease: low selenium status is common in Crohn’s disease and correlates inversely with the severity of experimental colitis (Speckmann & Steinbrenner, *Inflamm Bowel Dis* 2014), whereas higher selenium status has been associated with lower colorectal-cancer risk in prospective cohorts, most clearly in women (Hughes et al., *Int J Cancer* 2015), though interventional supplementation has not shown a consistent cancer-preventive benefit (Vinceti et al., *Cochrane Database Syst Rev* 2018). Yet how the intestinal epithelium takes up and deploys selenium, and how this is remodeled across inflammation and cancer, has remained poorly defined at cellular resolution.

Selenoprotein synthesis begins with the cellular uptake of selenium in a usable form. Dietary selenium reaches the gut lumen in several chemical species, among them selenocystine, the selenium analogue of cystine. We recently showed, in the healthy human intestine, that the apical brush-border system b^0,+^, the *SLC3A1* heavy chain (rBAT) and *SLC7A9* light chain (b⁰,⁺AT), whose loss causes cystinuria through failed luminal reabsorption of cystine and dibasic amino acids (Fotiadis et al., *Mol Aspects Med* 2013), is the membrane transport system most robustly associated with a selenoprotein-enriched transcriptional state in enterocytes, defining a selenium-utilization program (He et al., *bioRxiv* 2026). Because b^0,+^ imports both cystine and selenocystine, it supplies the substrates for glutathione synthesis and for selenoprotein (notably *GPX4*) production, placing it upstream of two convergent antioxidant outputs.

This positions b^0,+^ at the supply node of an epithelial redox axis with two opposing poles. A selenium pole runs b^0,+^ → selenocystine → the selenocysteine-incorporation machinery → *GPX4* and the wider selenoproteome; a thiol pole runs the cystine– glutamate antiporter system xc⁻ (xCT/*SLC7A11*) → cystine → glutathione, a program driven transcriptionally by NRF2 and frequently amplified in cancer (Koppula et al., *Protein Cell* 2021). Both poles converge on defence against ferroptosis, an iron-dependent form of regulated cell death caused by unrestrained phospholipid peroxidation (Dixon et al., *Cell* 2012; Jiang et al., *Nat Rev Mol Cell Biol* 2021), for which the selenoprotein GPX4 is the central guardian (Ingold et al., *Cell* 2018). Ferroptosis is implicated at both disease poles: it is induced in the intestinal epithelium in ulcerative colitis, contributing to epithelial cell death and barrier breakdown (Xu et al., *Cell Death Dis* 2020), and seleno-amino-acid supply protects the epithelial barrier through GPX4/glutathione and mitigates experimental IBD (Huangfu et al., *iScience* 2024); conversely, ferroptosis is a targetable dependency of therapy-resistant and drug-tolerant tumour cells, including in colorectal cancer (Hangauer et al., *Nature* 2017; Zhang et al., *Front Oncol* 2022). A single redox-currency axis could thus read out as vulnerability under inflammation and as resistance in cancer.

We therefore asked how intestinal disease remodels this axis: whether the loss of b^0,+^ is an isolated event or one node of a coordinated supply-defence program, and whether inflammation and cancer move the axis in opposite directions. Using a multi-cohort single-cell framework with donor-level statistics, cross-validated in bulk transcriptomic and proteomic cohorts, including an independent patient proteome and a Caco-2 polarization model, we find that inflammation and cancer remodel the axis in opposing directions, that the inflammatory loss of b^0,+^ reflects epithelial cell-state remodeling rather than transcriptional repression in differentiated cells, and that b^0,+^ marks the supply node of the axis; a direct isogenic test of whether removing b^0,+^ is sufficient to reproduce the disease-associated switch is outlined as the next step.

## Results

### R1 | A dual-pole epithelial redox axis is remodeled in opposite directions by inflammation and cancer

Having established the b^0,+^ selenium-utilization program in healthy enterocytes (He et al., *bioRxiv* 2026), we framed disease remodeling as a test of a single redox-currency axis with two ferroptosis-defensive poles, a selenium pole (b^0,+^ → selenocystine → *GPX4*/selenoproteome) and a thiol pole (xCT → cystine → glutathione), with b^0,+^ as its supply node (Fig 1A). We interrogated this axis across five layers of predominantly public, donor-level–controlled data, from a cross-disease discovery atlas to protein-level and in-vitro cross-validation (Fig 1B), beginning with the Pan-GI Atlas Extended (Oliver et al., *Nature* 2024; Fig 1C).

**Fig 1.**
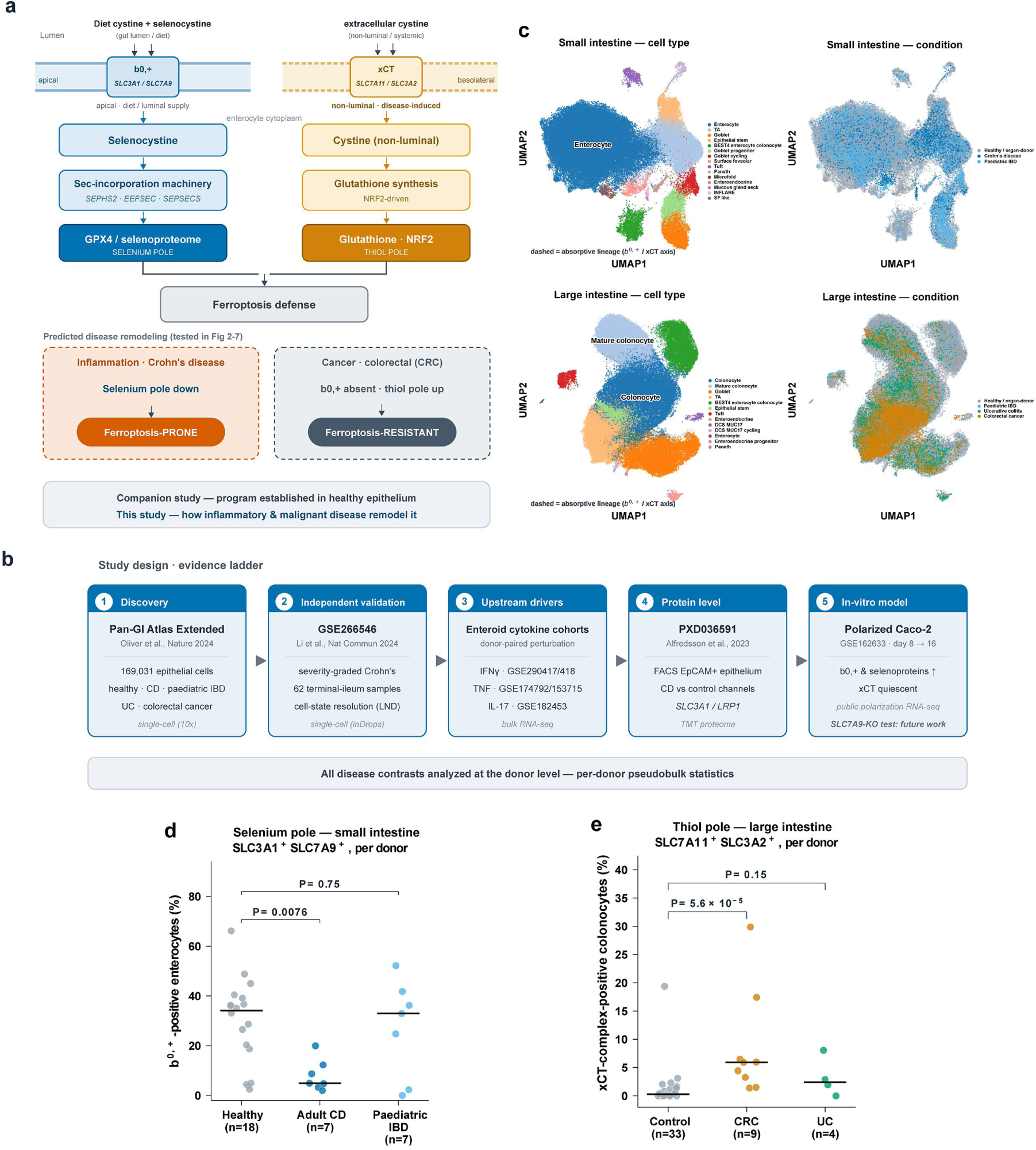
A dual-pole epithelial redox-currency axis is remodeled in opposite directions across inflammatory and malignant intestinal disease: framework, study design, and discovery. **(A)** Model under test. The b^0,+^ transporter (*SLC3A1* heavy chain / *SLC7A9* light chain) imports cystine and selenocystine at the enterocyte apical membrane, supplying two opposing, ferroptosis-defensive poles: a selenium pole (selenocystine → selenocysteine-incorporation machinery → *GPX4* and the wider selenoproteome) and a thiol pole (xCT/*SLC7A11* → cystine → glutathione under NRF2 control). Having established this program in the healthy human epithelium (He et al., bioRxiv 2026), we ask here how inflammatory and malignant disease remodel it. Predicted disease directions (inflammation collapses the selenium pole toward ferroptosis susceptibility; cancer bypasses b^0,+^ toward the thiol pole and ferroptosis resistance) are shown as hypotheses tested throughout. **(B)** Study design and evidence ladder. Five layers of predominantly public data, all analyzed with donor-level (per-donor pseudobulk) statistics: a cross-disease discovery atlas (Pan-GI Atlas Extended, Oliver et al., *Nature* 2024; 169,031 intestinal epithelial cells), an independent severity-graded Crohn’s-disease cohort (GSE266546; inDrops), upstream cytokine-perturbation small-intestinal enteroid cohorts (IFNγ: GSE290417/290418; TNF: GSE174792/GSE153715), an independent epithelial proteome (PXD036591), and a Caco-2 polarization model (GSE162633; b^0,+^/selenoprotein co-induction). All five layers are public transcriptomic/proteomic data; a direct isogenic *SLC7A9*-knockout functional test is outlined as future work**. (C)** UMAP of Pan-GI Atlas intestinal epithelial cells, colored by cell type (left) and by disease condition (right); small- and large-intestinal compartments indicated, locating the absorptive enterocytes and colonocytes profiled below. **(D)** Selenium-pole discovery. Donor-level b^0,+^ co-expression (% of small-intestinal enterocytes positive for both *SLC3A1* and *SLC7A9*; donors with ≥20 absorptive cells); each point one donor. All comparisons are made at the donor level (per-donor pseudobulk), not per cell, to avoid pseudoreplication (Squair et al., *Nat Commun* 2021). Healthy/organ-donor controls (n = 18) vs adult Crohn’s disease (n = 7): median 34.1% vs 4.9%, two-sided Mann–Whitney *P* = 0.008. Paediatric IBD (n = 7) does not differ from the same 18-donor healthy comparator (median 33.0%, *P* = 0.75), indicating the deficit is specific to adult, actively inflamed tissue rather than a generic IBD feature. **(E)** Thiol-pole discovery (contrast pole). Donor-level xCT-complex co-expression (% of large-intestinal colonocytes positive for both *SLC7A11* and *SLC3A2*; same threshold). Control (n = 33) vs colorectal cancer (n = 9): median 0.3% vs 5.9%, *P* = 5.6 × 10⁻⁵, by far the strongest single readout in the malignant epithelium. Ulcerative colitis (n = 4) shows an intermediate, non-significant trend (median 2.4%, *P* = 0.15), underpowered at this donor count. b^0,+^ is essentially absent from colonocytes in all conditions (Methods). The malignant pole is established here at the atlas/population level as the opposing configuration of the axis; its mechanistic dissection is beyond the scope of this study.

In the discovery atlas, the two poles moved in opposite directions. Small-intestinal enterocytes from adult Crohn’s disease donors showed a marked reduction in b^0,+^ co-expression (cells positive for both *SLC7A9* and *SLC3A1*): a per-donor median of 4.9% versus 34% in healthy donors (7 vs 18 donors, *P* = 0.008; continuous b^0,+^ score 0.28 vs 0.62, *P* = 0.034). The effect was specific to adult, actively inflamed tissue: paediatric IBD enterocytes showed no reduction relative to the same donor-level healthy comparator (33.0% vs 34.1%, *P* = 0.75) (Fig 1D). All significance testing throughout is performed at the donor level (per-donor pseudobulk), not per cell, to avoid the pseudoreplication that inflates significance when individual cells within a donor are treated as independent replicates; the reported *n* are therefore numbers of donors, the correct unit of biological replication (Squair et al., *Nat Commun* 2021).

The large-intestinal pole was the mirror image. b^0,+^ is essentially absent from colonocytes in all conditions, but colorectal cancer colonocytes strongly induced the xCT complex (*SLC7A11*+*SLC3A2* co-detection; median 5.9% vs 0.3% positive; 9 vs 33 donors, *P* = 5.6 × 10⁻⁵), by far the strongest single readout in the malignant epithelium; ulcerative colitis colonocytes showed an intermediate, non-significant trend (2.4% vs 0.3%; *n* = 4 donors, *P* = 0.15) (Fig 1E). We establish this malignant pole at the population level as the opposing configuration of the axis and return to it, together with its NRF2/ferroptosis-resistant character, in the integrative model (Fig 7); the mechanistic layers that follow (Fig 2–6) dissect the inflammatory pole, where the disease burden of this study lies. Profiling all nine candidate selenium-transport systems across the four dietary selenium forms (Fig S1) singled out the two selenocystine-transporting systems, b^0,+^ and xCT, as the coherent opposite-pole pair of this axis: b^0,+^ was the transporter suppressed in the inflamed small intestine and xCT the one induced in the colorectal-cancer colon. Transporters for the other three dietary forms also changed in disease, but along programs distinct from the selenocystine/selenoprotein axis, the selenomethionine carrier LAT1 was co-induced with xCT in cancer, and the differentiation-associated anion transporters DRA/PAT1 fell with dedifferentiation, reinforcing b^0,+^ and xCT as the selenoprotein-relevant supply nodes this study follows.

**Fig 2.**
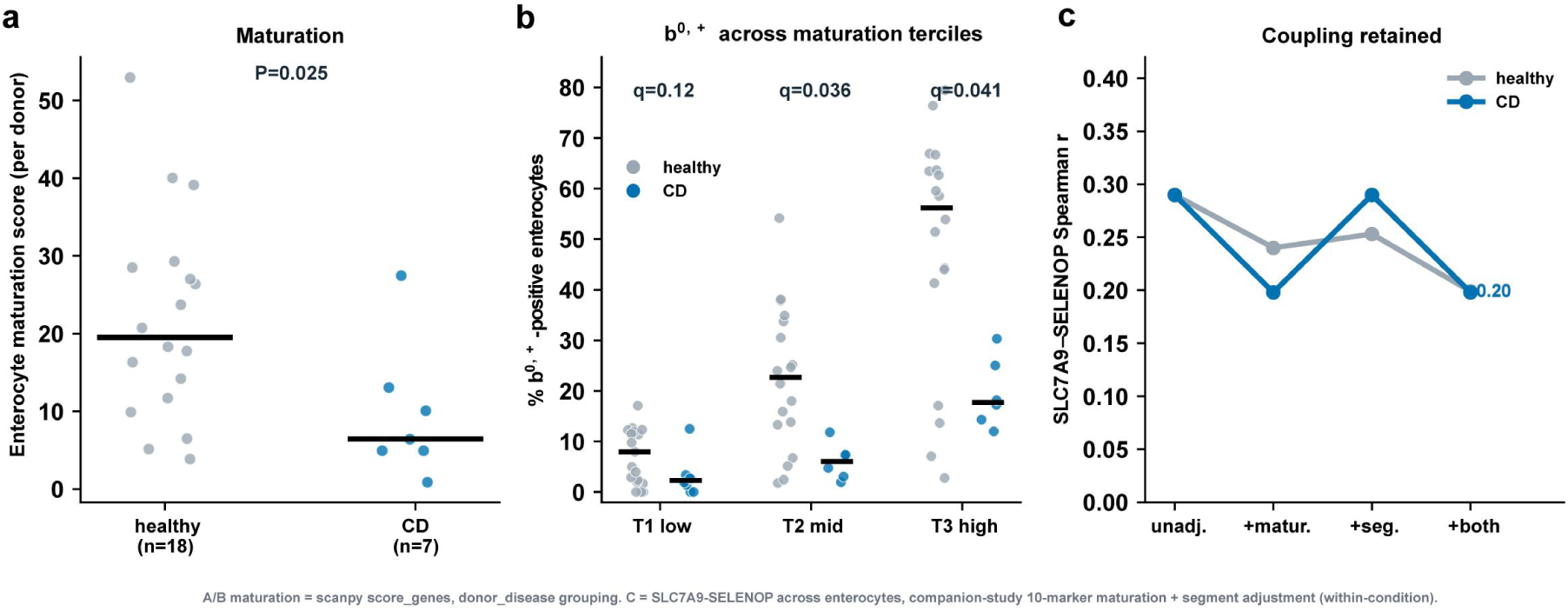
Loss of b^0,+^ in Crohn’s disease reflects depletion of the b^0,+^-high enterocyte state, not transcriptional disruption. Pan-GI Atlas Extended small-intestinal enterocytes, donor-level; healthy = organ-donor + control (n = 18) vs adult Crohn’s disease (n = 7); maturation scored by scanpy score_genes over villus-enterocyte markers (one definition throughout). **(A)** Per-donor enterocyte maturation, healthy vs CD (median 19.5 vs 6.4; two-sided Mann–Whitney *P* = 0.025). **(B)** b^0,+^-positive fraction across maturation terciles; the deficit persists in the most-mature tercile (56% vs 17.7% positive; BH *q* = 0.04), so it is not merely a consequence of reduced maturation. Each point is a per-donor b^0,+^-positive fraction *within* a maturation tercile (donors with ≥10 absorptive cells in that tercile); the test is performed at the donor level within each tercile. **(C)** *SLC7A9*–*SELENOP* coupling across enterocytes, reproducing the maturation-adjustment method of He et al. (bioRxiv 2026): the correlation attenuates as maturation and segment are adjusted for but is essentially identical in healthy and Crohn’s tissue (fully-adjusted partial Spearman *r* = 0.20 vs 0.20; ∼47% of the variance retained in both), i.e. the coupling is retained in disease. Together: quantitative depletion of the b^0,+^-high population with the program intact where retained.

**Fig 3.**
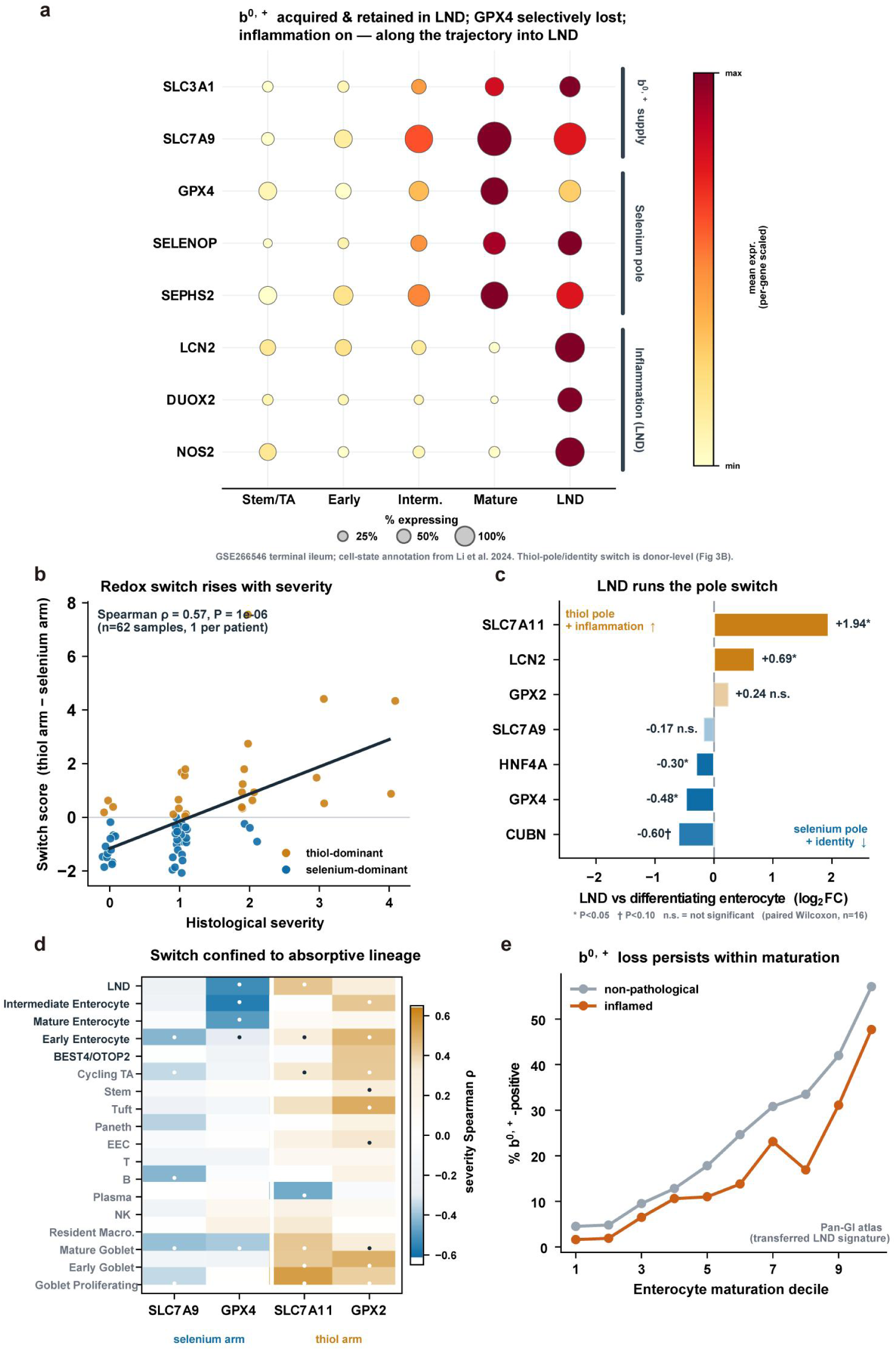
A dedifferentiated cell state (LND) carries the b^0,+^ loss and executes a lineage-restricted, severity-graded selenium→thiol pole switch. Severity-graded Crohn’s cohort (Vanderbilt Gut Cell Atlas, GSE266546; terminal ileum), with cross-cohort validation in the Pan-GI atlas. **(A)** Redox-axis genes across the absorptive-lineage trajectory (Stem/TA → Early → Intermediate → Mature → LND; GSE266546 terminal ileum, cell-state annotation from Li et al. 2024; dot size, % of cells expressing; colour, per-gene min–max-scaled mean expression). The b^0,+^ transporter (*SLC3A1*, *SLC7A9*) is acquired with maturation and *retained* in LND, the single-cell correlate of a compositional, not in-cell, loss of b^0,+^. *GPX4* is selectively reduced in LND while the wider selenoprotein module (*SELENOP*, *SEPHS2*) is preserved, and an inflammation program (*LCN2*, *DUOX2*, *NOS2*) switches on specifically in LND. The thiol-pole induction and enterocyte-identity loss are donor-level effects and are shown at pseudobulk in (B), not at single-cell resolution here. **(B)** Per-sample redox switch score (thiol arm − selenium arm; absorptive-lineage pseudobulk, LND included) rises with histological severity (Spearman *ρ* = 0.57, *P* = 1 × 10⁻⁶; n = 62 samples). **(C)** LND-versus-differentiating-enterocyte switch (donor-paired canonical log₂FC, n = 16; paired Wilcoxon): selenium pole and identity down (*GPX4* −0.48*, *HNF4A* −0.30*, cubilin *CUBN* −0.60†), thiol pole and inflammation up (*SLC7A11*/xCT +1.94*, *LCN2* +0.69*). The b^0,+^ subunit *SLC7A9* (−0.17, n.s.) and *GPX2* (+0.24, n.s.) are unchanged in this paired contrast, b^0,+^ loss reflects compositional replacement, not within-state repression (* *P* < 0.05, † *P* < 0.10). **(D)** Cell-type × gene severity-correlation heatmap: selenium-arm suppression (*SLC7A9*/*GPX4*, blue) and thiol-arm induction (*SLC7A11*/*GPX2*, orange) are confined to the absorptive lineage (including LND) and mature goblet cells, and absent from progenitor, immune and stromal compartments (within-cell-type *ρ* = −0.33 to−0.60 for *GPX4*). **(E)** Cross-cohort validation: a curated LND signature transferred into the Pan-GI atlas (b^0,+^/selenoprotein/xCT readout genes excluded to avoid circularity) reproduces lower b^0,+^-positive fractions in inflamed than non-pathological regions across all enterocyte maturation deciles.

**Fig 4.**
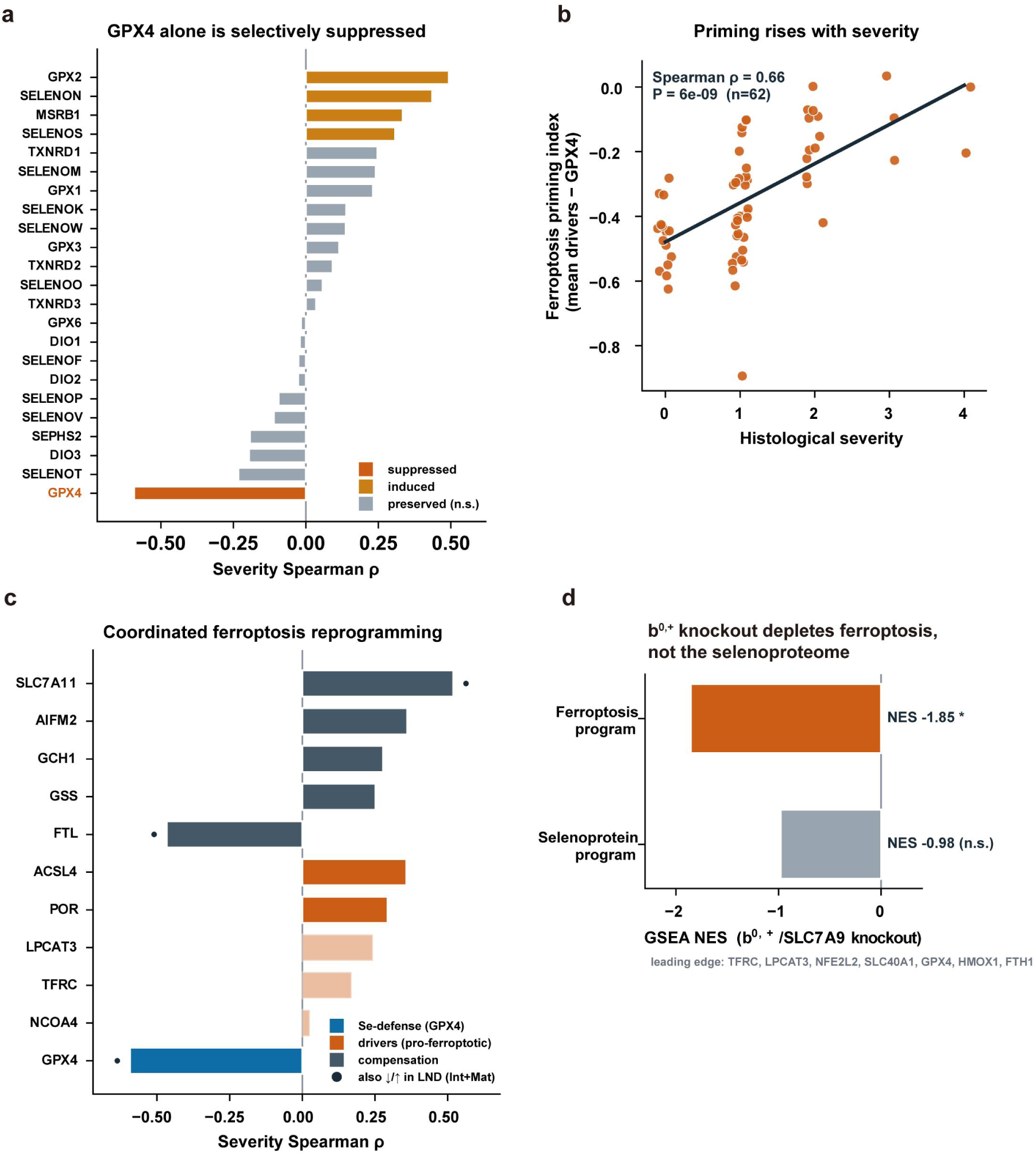
A selective loss of GPX4 primes the inflamed epithelium for ferroptosis. Severity-graded Crohn’s cohort (GSE266546, terminal ileum; absorptive-lineage pseudobulk) plus in-silico network knockout. **(A)** Across 23 detected selenoproteins ranked by severity correlation, *GPX4* alone is selectively suppressed (Spearman ρ = −0.59); the rest are preserved or induced and the selenocysteine-incorporation machinery (*SEPHS2* shown; *SECISBP2*, *EEFSEC*, *PSTK*, *SEPSECS*) is unchanged, a *GPX4*-selective loss on preserved incorporation machinery (substrate-supply versus *GPX4*-intrinsic mechanisms discriminated by the functional test in the Discussion). **(B)** A ferroptosis priming index (per-sample mean of pro-ferroptotic drivers minus *GPX4*) rises with histological severity (Spearman ρ ≈ +0.66). **(C)** The ferroptosis program reprograms coordinately with severity, the selenium-defense guardian *GPX4* falls (blue), pro-ferroptotic drivers rise (*ACSL4*, *POR*; vermillion) and compensation genes rise (xCT, *AIFM2*/FSP1, *GCH1*; slate) with ferritin *FTL* lost (releasing labile iron), leaving a net susceptibility window; filled dots mark genes also significant in the LND-versus-differentiating-enterocyte contrast (Intermediate + Mature reference, matching Fig 3). **(D)** In-silico network knockout of the b^0,+^ catalytic subunit *SLC7A9* depletes the ferroptosis program (GSEA NES = −1.85, *P* = 0.017; leading edge *GPX4*, *FTH1*, *SLC40A1*, *TFRC*) but not the selenoprotein program (NES = −0.98, n.s.), a functional supply→vulnerability link rather than a direct transcriptional b^0,+^→selenoprotein axis (see also Fig S3).

**Fig 5.**
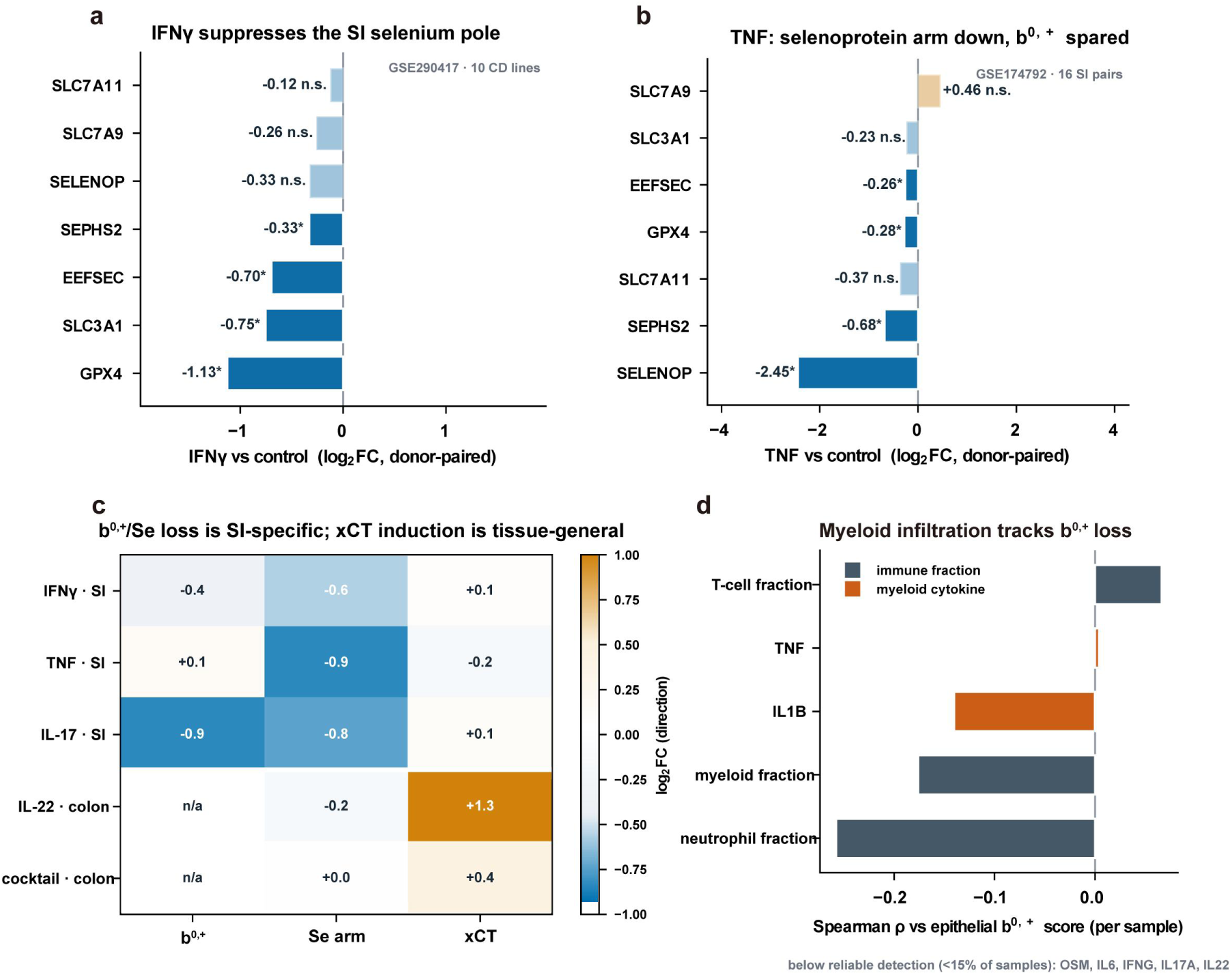
Inflammatory cytokines suppress the small-intestinal selenium pole (bulk enteroids), which b^0,+^ supports through substrate supply rather than transcription. Public small-intestinal enteroid cytokine RNA-seq (single-laboratory; donor-paired) plus the severity-graded tissue cohort. **(A)** IFNγ, in ten donor-paired Crohn’s-disease enteroid lines, lowers the guardian *GPX4* (log₂FC = −1.13), the b^0,+^ heavy chain *SLC3A1* (−0.75) and the selenocysteine machinery *SEPHS2*/*EEFSEC* (all BH *q* ≤ 0.017); *SLC7A9*, *SELENOP* and xCT are unchanged (GSE290417; concordant, underpowered in six non-IBD lines, GSE290418). **(B)** TNF, in two paired small-intestinal datasets (GSE174792, n = 16 pairs shown), suppresses the selenoprotein effectors and machinery (*SELENOP*−2.45, *SEPHS2* −0.68, *GPX4* −0.28, *EEFSEC* −0.26; * paired Wilcoxon *P* < 0.05) while leaving the b^0,+^ transporter unchanged (*SLC3A1*, *SLC7A9*, n.s.), the two arms are independently regulated. **(C)** Directional summary across verified datasets: b^0,+^/selenoprotein-pole suppression requires a small-intestinal model and a pro-inflammatory cytokine, whereas xCT induction is tissue-general (seen in colonic models, e.g. IL-22; b^0,+^ “n/a” where not expressed). **(D)** In the severity-graded tissue (GSE266546), per-sample Spearman correlation of immune-cell fractions and detectable cytokines against the epithelial b^0,+^ score: myeloid and neutrophil infiltration and IL1B track b^0,+^ loss, while T-cell cytokines (IFNG, IL17A, IL22) fall below reliable detection (< 15% of samples).

**Fig 6.**
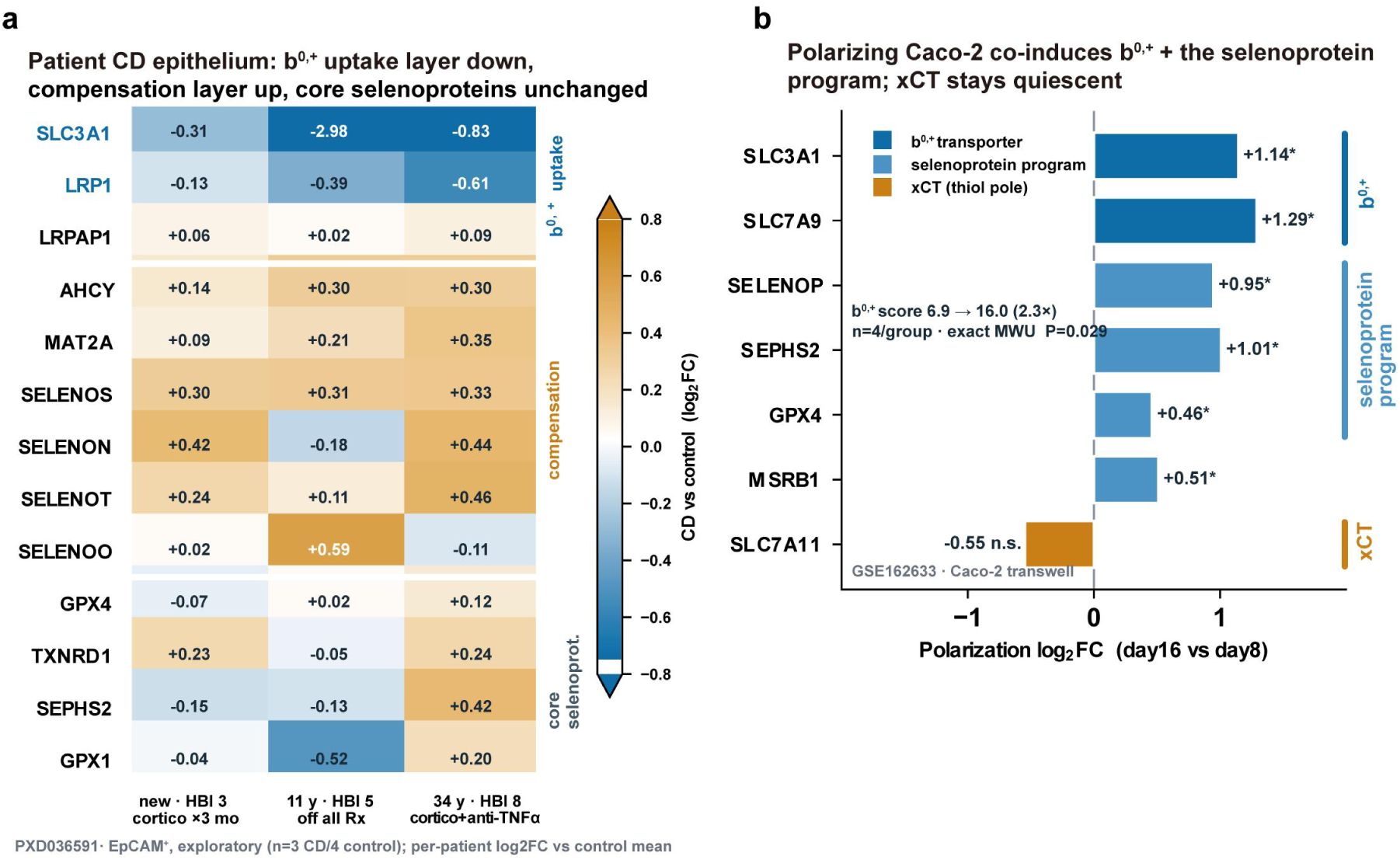
Protein-level and cellular-model cross-validation of the b^0,+^ supply node. **(A)** Motivating public patient proteome (PXD036591; exploratory, *n* = 3 CD / 4 control EpCAM⁺ channels), per-patient log₂FC (CD channel vs control mean) shown as a two-layer heatmap, patients ordered by disease duration/HBI and annotated by current treatment (PXD036591 clinical metadata, Alfredsson et al.). The b^0,+^ uptake layer is reduced in every patient: *SLC3A1* pooled log₂FC −1.01 (per-patient −0.31/−2.98/−0.83, largest in the intermediate-duration patient), its endocytic partner *LRP1* −0.36 (−0.13/−0.39/−0.61), while the *LRP1* chaperone *LRPAP1* is unchanged (specificity control). A methionine-cycle/ER-stress compensation layer is coordinately, modestly up (*AHCY*, *MAT2A*, *SELENOS*, *SELENOT* up in all three patients; *SELENON*/*SELENOO* variable), whereas core selenoproteins (*GPX4*, *TXNRD1*, *SEPHS2*, *GPX1*) are unchanged, localizing the deficit to the b^0,+^ supply/uptake node rather than the selenoprotein machinery. Direction only (small n, not powered for significance). **(B)** Model validation in independent public data (GSE162633): as Caco-2 polarizes on Transwell (day 8 → day 16, *n* = 4 per timepoint), the b^0,+^ transporter (*SLC3A1* +1.14, *SLC7A9* +1.29 log₂FC) and the selenoprotein-utilization program (*SELENOP* +0.95, *SEPHS2* +1.01, *MSRB1* +0.51, *GPX4* +0.46; all exact Mann–Whitney *P* = 0.029) are co-induced, while the thiol-pole transporter xCT (*SLC7A11*) is not (−0.55, n.s.), the polarized monolayer runs the selenium pole with the thiol pole quiescent, the intended background for perturbing b^0,+^. Effect direction/magnitude only (two timepoints, *n* = 4); *P* = 0.029 is the exact Mann– Whitney floor at this sample size.

**Fig 7.**
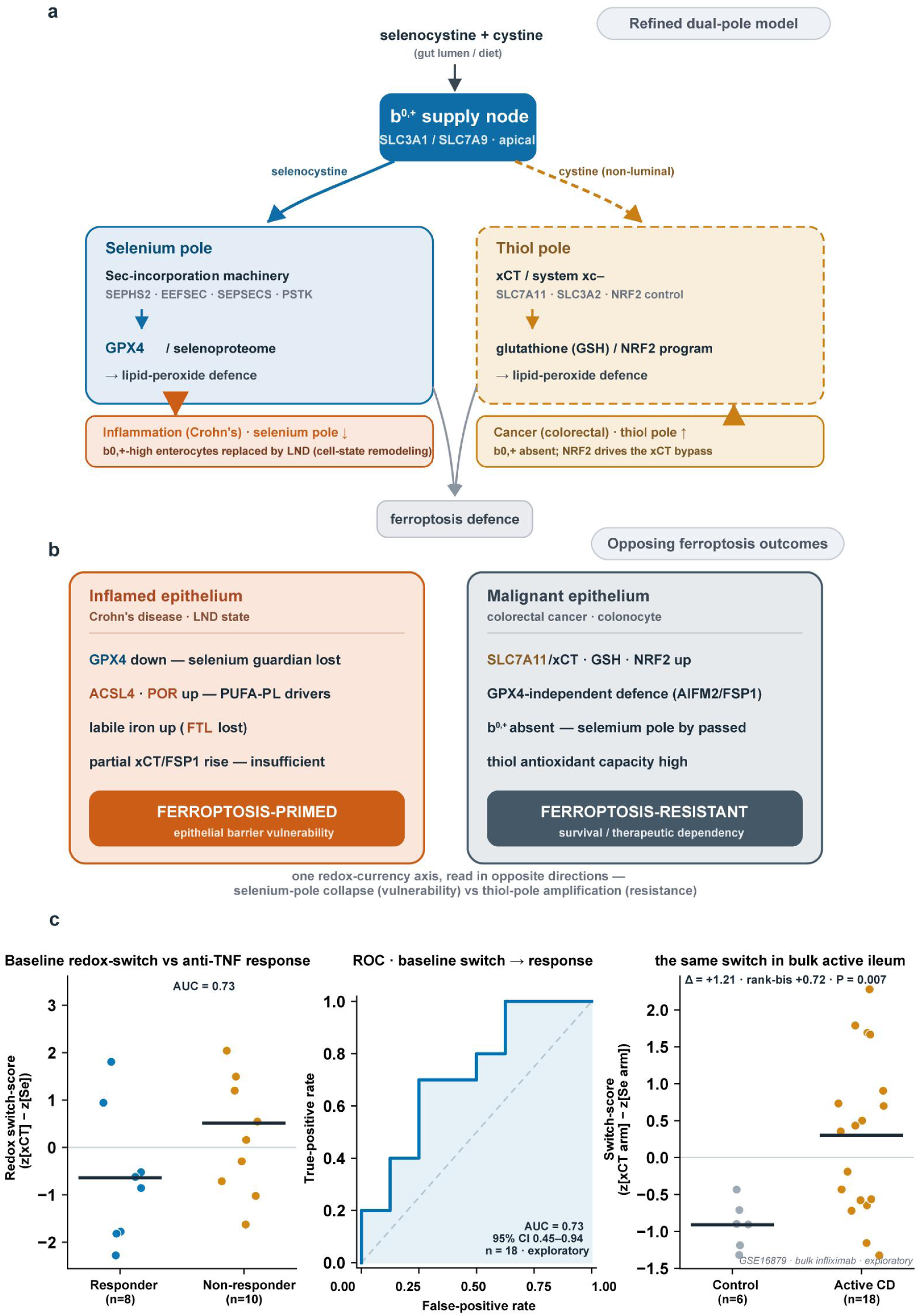
Integrative dual-pole redox-currency model, the malignant contrast pole, and an exploratory anti-TNF baseline correlate. **(A)** Refined dual-pole model (validated counterpart of the Fig 1A hypothesis): b^0,+^ supply node feeding selenium and thiol poles, with inflammation suppressing the selenium pole via cell-state remodeling (Fig 2–4) and cancer bypassing it toward the NRF2/xCT thiol pole. **(B)** Opposing ferroptosis outcomes: ferroptosis-primed inflamed epithelium vs ferroptosis-resistant malignant epithelium. **(C)** Exploratory clinical correlate in GSE16879: (left) baseline composite switch-score in infliximab responders vs non-responders (*n* = 8 vs 10); (middle) ROC of the baseline-switch classifier (AUC = 0.73; bootstrap 95% CI 0.45–0.94, spanning 0.5, hypothesis-generating, not a validated biomarker); (right) the switch reproduced in bulk active Crohn’s ileum (Δ = +1.21, rank-biserial +0.72, *P* = 0.007).

### R2 | Loss of b^0,+^ in Crohn’s disease reflects depletion of the b^0,+^-high enterocyte state, not transcriptional disruption

How is the b^0,+^ program lost? Two explanations differ in what is affected: the program could be disrupted within enterocytes that still express it, or the b^0,+^-high cell state could be depleted from the epithelium while the program stays intact in the cells that retain it. A single-cell atlas separates these, because it reads both the per-cell program and cell-state composition; what it cannot say is how the state is depleted, i.e., whether mature cells lose it by dedifferentiation, die, or fail to acquire it through a maturation block. Two observations favour depletion of the b^0,+^-high state over in-cell disruption. Crohn’s enterocytes were less mature than control enterocytes (per-donor maturation score 6.4 vs 19.5, *P* = 0.025; Fig 2A), and the b^0,+^ deficit persisted even at matched maturity: 17.7% vs 56% positive in the most-mature tercile (Fig 2B), so it is not merely a consequence of the maturation shift. A reduced detection rate at matched maturity could in principle reflect either fewer b^0,+^-high cells within the stratum or lower b^0,+^ within the cells that remain; the *SLC7A9*–*SELENOP* coupling distinguishes these. This coupling, which represent the strongest maturation-independent b^0,+^ association established in healthy tissue (He et al., *bioRxiv* 2026), was retained across Crohn’s enterocytes: after adjusting for maturation and intestinal segment it was essentially unchanged (partial *r* = 0.20 in Crohn’s vs 0.20 in controls, ∼47% of the association’s variance surviving adjustment in both groups; Fig 2C), and specific to SELENOP as in the companion study. The pseudobulk *SLC7A9*–*SELENOP* relationship likewise remained strong.

Together these indicate that the b^0,+^-high enterocyte population is quantitatively depleted and replaced by less mature, b^0,+^-low cells, rather than the program being switched off within differentiated enterocytes, via depletion rather than disruption.

### R3 | A dedifferentiated cell state (LND) carries the b^0,+^ loss and executes the selenium-to-thiol pole switch

R2 established that Crohn’s disease depletes the b^0,+^-high enterocyte and enriches for less-mature cells with the program intact where retained, but the mixed-severity Pan-GI atlas cannot name the cell state that replaces the b^0,+^-high enterocyte or place it on a severity axis. We therefore turned to an independent, histologically severity-graded Crohn’s cohort with single-cell resolution of dedifferentiated epithelium (GSE266546; Li et al., *Nat Commun* 2024). There, the remodeling resolved into a discrete inflammation-associated state, large non-differentiated (LND) cells, that expanded with disease and replaced mature enterocytes. Across the absorptive-lineage trajectory (Stem/TA → mature enterocyte → LND), the b^0,+^ transporter (*SLC3A1*, *SLC7A9*) was acquired with differentiation and retained in LND, while *GPX4* declined, whereas the wider selenoprotein module (*SELENOP*, *SEPHS2*) preserved, and an inflammation program (*LCN2*, *DUOX2*, *NOS2*) switched on specifically in LND (Fig 3A). Across samples, this redox switch was graded by disease, whereby the per-sample switch score (thiol arm − selenium arm) rose with histological severity (Spearman *ρ* = 0.57, *P* = 1 × 10⁻⁶; Fig 3B). Decomposed at the cell-state level, LND cells executed the switch relative to differentiating enterocytes (donor-paired, n = 16; Fig 3C): suppression of the selenium pole (*GPX4* log₂FC = −0.48, *P* = 0.02) and of enterocyte identity (*HNF4A* −0.30, *P* = 0.001; apical endocytic receptor cubilin *CUBN* −0.60, *P* = 0.06) with induction of the thiol pole and inflammation (*SLC7A11*/xCT +1.94, *P* = 0.008; *LCN2* +0.69, *P* < 10⁻³). Critically, the b^0,+^ catalytic subunit *SLC7A9* was not itself transcriptionally lower in LND (−0.17, *P* = 0.13), so, consistent with the depletion-not-disruption finding of R2, the tissue-level b^0,+^ deficit reflects compositional replacement of mature enterocytes by the low-b^0,+^ LND state rather than in-cell repression of the transporter; the switch LND executes is one of the downstream redox effectors (*GPX4*, xCT), not of the b^0,+^ supply arm. *GPX2*, by contrast, was not uniformly induced across the differentiating pool (+0.24, non-significant) but tracked severity within the absorptive lineage (Fig 3D).

Two further features confirmed LND as a genuinely dedifferentiated, non-proliferative state rather than an expanding progenitor. First, LND was the least proliferative epithelial state, not the most: cell-cycle scoring placed only 8.6% of LND cells in S/G2M, a proportion comparable to mature enterocytes (9.7%) and far below the stem/transit-amplifying positive control (43.5%). In a donor-paired comparison within active disease, LND cells exhibited lower cycling rates than mature enterocytes (median 7.0% vs 10.9%, Wilcoxon *P* = 0.007, n = 12 donors). This was not a depth artifact, as LND carried the highest per-cell sequencing depth of any state (Fig S2A–C). LND accumulation therefore reflects dedifferentiation of existing epithelium, not clonal expansion of a proliferative progenitor. Second, the identity loss was coordinated at the gene-set level rather than driven by individual markers: the mature-enterocyte identity program was enriched for down-regulation both along the LND-versus-mature axis (prerank GSEA NES = −1.92, FDR = 0.009) and along the independent histological-severity axis (NES = −2.38, FDR = 0.013; k = 19 leading-edge genes including *HNF4A*, *SI*, *ALPI*, *FABP1*, *MTTP*; Fig S2D). A batch-corrected trajectory analysis (Harmony integration over sample of origin) placed LND at the differentiated end of the absorptive lineage, with the strongest connectivity to intermediate and mature enterocytes (PAGA connectivity 0.63 and 0.79) and least to stem cells (0.09), and diffusion pseudotime values adjacent to those of mature enterocytes (Fig S2E). This positioning supports LND as a a dedifferentiated offshoot of maturing enterocytes rather than a stem-like state, and this topological feature remains stable across uncorrected, ComBat and Harmony batch treatments.

This xCT induction is confined to the dedifferentiated LND compartment and emerges with severity: in the mature small-intestinal enterocytes profiled in the mixed-severity Pan-GI atlas, xCT remains low and is not significantly altered in Crohn’s disease (Fig S1), so the apparent atlas-versus-cohort discrepancy resolves once cell state and severity are accounted for.

This switch was confined to the differentiated epithelial compartment. Severity-associated *GPX4* suppression occurred only in absorptive enterocytes, LND and mature goblet cells (within-cell-type ρ = −0.33 to −0.60) and was absent from progenitor, immune and stromal compartments; the b^0,+^ substrate arm (*SLC7A9*, *SLC3A1*) was expressed almost exclusively in absorptive cells. Within mature enterocytes, *SLC7A9* did not itself fall with severity (ρ = −0.07), consistent with the tissue-level loss arising from the shift in composition from mature enterocytes to the LND state. To test whether this cell-state basis generalized beyond the severity-graded cohort, we turned to the predominantly healthy Pan-GI atlas, which contributes cross-disease breadth where GSE266546 contributes cell-state resolution. Because the raw LND markers index well-differentiated epithelium in this largely healthy tissue, we instead transferred a curated maturation/inflammation signature derived from the GSE266546 LND-versus-mature contrast (excluding b^0,+^, selenoprotein and xCT readout genes to avoid circularity) into the atlas’s small-intestinal enterocytes. The resulting LND-high subpopulation expanded specifically in inflamed sample regions, carried lower b^0,+^ and *GPX4* than LND-low cells, and retained b^0,+^ suppression within maturation deciles. These findings recapitulate, in an independent atlas, the cell-state basis established directly in GSE266546 (Fig 3E).

### R4 | Downstream: a selective loss of *GPX4* primes the inflamed epithelium for ferroptosis

The redox polarity shift executed within LND cells disrupts the epithelial selenium-defence axis; its functional cost depends on which selenoproteins become limiting. Profiling the selenoproteome, we found *GPX4* to be the only selenoprotein selectively suppressed with severity (Spearman ρ = −0.59, ranked first of 23 detected; Fig 4A), and the priority hierarchy expected under systemic selenium deficiency was absent; the selenocysteine-incorporation machinery (*SEPHS1/2*, *SECISBP2*, *EEFSEC*, *PSTK*, *SEPSECS*) was preserved. This selectivity argues against a global collapse of selenoprotein synthesis and does not match the priority-graded pattern of systemic selenium deficiency; it points to a *GPX4*-specific lesion. At the transcript level, this *GPX4*-selective decline is part of the LND redox-and-identity switch, whereby *GPX4* falls with enterocyte-identity genes (*HNF4A*, *CUBN*; R3) rather than in isolation. This transcriptional co-regulation and b^0,+^-dependent supply are not mutually exclusive but act at different levels: b^0,+^ sets the selenocysteine available to whatever *GPX4* is transcribed, so a supply deficit, with a possible *GPX4*-intrinsic, oxidant-sensitive translational component, would compound the transcriptional decline rather than replace it. Which contribution dominates the correlative data cannot settle; the isogenic *SLC7A9* knockout outlined in the Discussion is designed to resolve it, testing whether removing b^0,+^ lowers *GPX4* output independently of identity loss.

Losing the epithelium’s principal lipid-peroxide guardian left the differentiated compartment ferroptosis-primed, and priming scaled with disease: a composite index contrasting the pro-ferroptotic execution arm against *GPX4* rose steeply with histological severity (Spearman ρ = +0.66; Fig 4B). The reprogramming was coordinated across the three ferroptosis arms (Fig 4C): within the LND state, relative to differentiating enterocytes, the Intermediate + Mature reference of Fig 3, *GPX4* fell (log₂FC = −0.48, *P* = 0.02) alongside the iron-storage ferritin *FTL* (−0.39, *P* < 10⁻³), releasing labile iron; the pro-ferroptotic drivers trended upward without reaching individual significance in this paired contrast (*ACSL4* +0.41, non-significant); and the coincident induction of xCT (+1.94) marked a partial, thiol-based compensation rather than rescue, leaving a net susceptibility window. That the vulnerability is anchored to the supply node itself was confirmed in silico (scTenifoldKnk; Osorio et al., *Patterns* 2022): network knockout of the b^0,+^ catalytic subunit *SLC7A9* depleted the ferroptosis program (GSEA NES = −1.85, *P* = 0.017) while leaving the selenoprotein program intact (Fig 4D), representing a functional supply→vulnerability link, not a direct transcriptional b^0,+^→selenoprotein axis. What severs that supply upstream, we turn to next.

### R5 | Upstream: inflammatory cytokines suppress the small-intestinal selenium pole, which b^0,+^ supports through substrate supply rather than transcription

To identify inflammatory signals upstream of the selenium pole, we screened public bulk-transcriptomic datasets of intestinal epithelium under cytokine or disease perturbation (86 human RNA-seq series retrieved; Methods). Being bulk transcriptional readouts of static enteroid cultures, these data establish the direction of an upstream driver but capture neither the dynamic epithelial turnover and cell-state redistribution that govern tissue-level b^0,+^ abundance in Crohn’s disease nor the single-cell resolution of the tissue analyses above; they therefore do not overturn the resolved in-vivo picture (R2– R4). The qualifying datasets that directly interrogated the b^0,+^/selenoprotein program were small-intestinal enteroid cytokine experiments, which in current public data come predominantly from one group (Sung Noh Hong and colleagues, Samsung Medical Center, South Korea; experimentally and temporally distinct studies and sample sets). There, IFNγ suppressed the selenium pole across ten donor-paired Crohn’s-disease-derived enteroid lines, the guardian *GPX4* (log₂FC = −1.13, BH q = 0.007) with the b^0,+^ heavy chain *SLC3A1* (−0.75, q = 0.007) and, in this bulk readout, the selenocysteine machinery *SEPHS2* (−0.33, q = 0.009) and *EEFSEC* (−0.70, q = 0.017; GSE290417, Lee et al., *Front Immunol* 2025; Fig 5A). Concordant but underpowered suppression was observed in six non-IBD lines from the same study (GSE290418); the b^0,+^ catalytic subunit *SLC7A9*, the composite b^0,+^ score and *SELENOP* did not reach significance (q = 0.43, 0.09, 0.36). That IFNγ lowers *SLC3A1* transcriptionally in this acute bulk model does not make transcriptional repression the dominant in-vivo route to b^0,+^ loss, as the single-cell tissue data attributes that loss to compositional replacement (R2). The cytokine effects that generalize across models (detailed below) act on the selenoprotein arm, not the transporter. Between the two cohorts, fold-change magnitudes did not differ (Mann–Whitney P = 0.26–0.96), so no disease-specific IFNγ hypersensitivity is claimed.

Two further paired datasets from the same group tested TNF directly in small-intestinal enteroids (GSE174792 and GSE153715; n = 16 and n = 12 donor pairs; Fig 5B). In both, TNF converged on the same selenium pole through a *different node*: it suppressed the selenoprotein effectors and machinery (*GPX4*, *SELENOP*, *SEPHS2*, *EEFSEC*; paired Wilcoxon *P* < 0.05 in both datasets) while leaving the b^0,+^ transporter itself unchanged (*SLC3A1*/*SLC7A9*, non-significant). That TNF can collapse the selenoprotein arm without touching b^0,+^ shows the two arms are independently regulated, reinforcing that the b^0,+^ transporter is not the transcriptional driver of the selenoproteome (below) and that its disease-associated loss is compositional (R2–R3), not cytokine-imposed. IL-17 (GSE182453, an unpublished small-intestinal deposit) gave a directional but line-variable signal, interpretable only in the one line with a robust b^0,+^ baseline, and is reported as consistent rather than confirmatory; a mouse organoid IFNγ dataset (GSE282271) was excluded (single pooled line; different species).

Two limitations bound this inference. First, the small-intestinal cytokine evidence, though drawn from experimentally and temporally distinct studies, derives largely from a single laboratory’s enteroid system; independent cross-laboratory replication in small intestine is lacking, because small-intestinal cytokine-perturbation datasets are rare. Second, the one response that reproduced across independent (colonic) laboratories was induction of xCT, a tissue-general thiol-pole response that mirrors the selenium→thiol switch of R3, rather than suppression of the small-intestinal b^0,+^/selenoprotein pole (Fig 5C). Consistent with an inflammatory origin, the epithelial-cytokine sources detectable in the severity-graded tissue were innate myeloid cells (IL1B, TNF and OSM in recruited macrophages and neutrophils); myeloid and neutrophil infiltration and the IL1B signal tracked the epithelial b^0,+^ loss (Fig 5D), while T-cell cytokines, the source of IFNγ, fell below detection of the inDrops platform.

Finally, does inflammation suppress b^0,+^ and the selenoproteome through a direct b^0,+^→selenoprotein transcriptional link, or by independently targeting each? Network-based virtual knockout (scTenifoldKnk; Osorio et al., *Patterns* 2022) in the physiological small-intestinal enterocyte network favours independent targeting: deleting either b^0,+^ subunit produced no depletion of the selenoprotein gene set (GSEA *P* ≥ 0.49 for both *SLC3A1* and *SLC7A9*; Fig S3), whereas knockout of the catalytic subunit *SLC7A9*, but not the heavy chain *SLC3A1*, depleted the ferroptosis program (NES = −1.85; the supply→vulnerability link already established in R4, Fig 4D). b^0,+^ therefore supports the selenoproteome post-transcriptionally, most simply via selenium-substrate supply rather than transcriptional control, exactly as TNF’s sparing of the b^0,+^ transporter predicts. Because a co-expression network cannot model substrate supply, protein translation or ferroptosis execution, this prediction ultimately requires direct functional testing, which we outline as a proposed isogenic-knockout experiment in the Discussion.

Two orthogonal public datasets corroborate this supply-node model beyond single-cell transcriptomics. At the protein level, we re-analysed a public isobaric-labelling (TMT) proteome of FACS-purified EpCAM⁺ epithelium from Crohn’s-disease and control biopsies (PXD036591; Alfredsson et al., *Proteomics* 2023; *n* = 3 Crohn’s-disease and 4 control channels, exploratory). The b^0,+^ uptake layer was reduced in every patient: *SLC3A1* pooled log₂FC −1.01 (per-patient −0.31, −0.83 and −2.98) and its endocytic partner *LRP1* −0.36 (−0.13, −0.39, −0.61). Meanwhile the *LRP1* chaperone *LRPAP1* and core selenoproteins (*GPX1/2/4*, *TXNRD1/2*, *SEPHS2*) showed no consistent change and a methionine-cycle/ER-stress compensation layer (*AHCY*, *MAT2A*, *SELENOS*, *SELENOT*) was modestly, coordinately regulated, revealing a two-layer response in which the b^0,+^ supply node is selectively lost at the protein level while the epithelium mounts a compensatory sulfur/selenium-handling program (Fig 6A). The magnitude of loss tracked treatment rather than disease duration or activity index. The largest *SLC3A1* reduction (−2.98) was in the only patient off all therapy, whereas the two on corticosteroids ± anti-TNFα showed much smaller reductions (PXD036591 clinical metadata, Alfredsson et al.). This pattern supports an inflammation-driven, potentially treatment-reversible loss of b^0,+^ transport. Though with three patients and mixed regimens this is a directionally concordant observation rather than confirmatory evidence, and treatment is itself a confounder of the disease-versus-control contrast.

In vitro, the same coupling is recapitulated in an isogenic system. As Caco-2 cells polarise to a small-intestinal absorptive-enterocyte phenotype (GSE162633; day 8 → day 16, *n* = 4 per timepoint), the b^0,+^ transporter (*SLC3A1* +1.14, *SLC7A9* +1.29 log₂FC) and the selenoprotein-utilisation program (*SELENOP* +0.95, *SEPHS2* +1.01, *MSRB1* +0.51, *GPX4* +0.46; all exact Mann–Whitney *P* = 0.029, the floor at this sample size) are co-induced. In contrast, the thiol-pole transporter xCT (*SLC7A11*) remains unchanged (−0.55, n.s.). This differentiated monolayer operates the selenium pole while the thiol pole remains quiescent, and thus serves as the natural background for a direct functional test of b^0,+^ loss, which we outline as the immediate next step (Discussion; Fig 6B).

### R6 | An integrative dual-pole model, the malignant contrast pole, and a candidate treatment-response correlate

Together, the preceding sections resolve a single epithelial redox-currency axis and how disease remodels it. b^0,+^ is its supply node, importing selenocystine and cystine at the apical membrane (He et al., *bioRxiv* 2026) and feeding two opposing, ferroptosis-defensive poles: a selenium pole (selenocystine → selenocysteine-incorporation machinery → *GPX4* and the wider selenoproteome) and a thiol pole (xCT → cystine → glutathione under NRF2 control). Inflammation and cancer drive the axis in opposite directions. Inflammation collapses the selenium pole through epithelial cell-state remodeling (R2–R4), while cancer bypasses b^0,+^ toward the thiol pole, yielding the refined, evidence-anchored model of Fig 7A, the validated counterpart of the Fig 1A hypothesis.

The malignant pole is the mirror image of the inflammatory one. Colorectal-cancer colonocytes induced the xCT complex at the donor level (*SLC7A11*/*SLC3A2*; median 0.3% vs 5.9% positive, *P* = 5.6 × 10⁻⁵; Fig 1E) on a background of near-absent b^0,+^, adopting an NRF2-associated, ferroptosis-resistant configuration, which stands in the opposite of the ferroptosis-primed inflamed epithelium of R4. The same axis therefore reads out as vulnerability under inflammation and as resistance in cancer (Fig 7B). We establish the malignant pole here at the atlas/population level, to complete the axis rather than to dissect it: a cell-state, severity and functional analysis of the cancer epithelium, analogous to the one applied to the inflammatory pole, is a distinct study beyond the present scope.

Finally, anchored on the inflammatory pole, we asked whether the baseline switch state carries prognostic information. In pre-treatment bulk cohorts (Arijs et al., *PLoS One* 2009; GSE16879), Crohn’s-disease patients who subsequently responded to infliximab had a lower baseline switch-score, corresponding to higher b^0,+^/selenium and lower xCT levels, than non-responders (composite switch-score AUC = 0.73, bootstrap 95% CI 0.45–0.94; *n* = 8 vs 10; Fig 7C), and the switch itself reproduced as an active-disease-versus-control contrast in bulk ileum (Δ = +1.21, rank-biserial +0.72, *P* = 0.007; *n* = 18 vs 6). Given the small samples and the bulk platform, this is a hypothesis-generating association rather than a validated biomarker; but it suggests that the same supply-node axis that grades disease severity may also track treatment response, a possibility we take up in the Discussion.

## Discussion

b^0,+^ is not an isolated transporter but the supply node where three programs intersect: nutrient supply (b^0,+^ importing cystine/selenocystine), redox and ferroptosis defense (the downstream glutathione and selenoprotein outputs), and the inflammatory/nutrient-sensing environment that sets whether the program is maintained. Our central finding is that intestinal disease remodels this axis as a coordinated unit and in disease-specific directions: Crohn’s disease suppresses the selenium pole, colorectal cancer bypasses it toward the thiol pole.

The mechanism of inflammatory suppression is cell-state remodeling. b^0,+^-high mature enterocytes are replaced by a dedifferentiated, ferroptosis-primed state (LND) rather than silenced in situ; the *SLC7A9*–*SELENOP* coupling is retained across disease enterocytes, and the loss is graded by severity and confined to the differentiated absorptive lineage. This reconciles the quantitative depletion seen across cohorts with the preservation of program quality, and integrates the otherwise competing possibilities of active suppression and passive dedifferentiation into a single remodeling model. The selective loss of *GPX4* on a preserved incorporation machinery, with concurrent *ACSL4* and xCT induction, renders the inflamed epithelium ferroptosis-prone, a cellular state mechanistically linked in broader literature to the barrier breakdown that characterizes active IBD.

The cancer pole is the inverse: with b^0,+^ absent, malignant colonocytes meet their cystine demand through xCT and an NRF2-driven thiol program, yielding ferroptosis resistance. The same axis therefore reads out as vulnerability in inflammation and as resistance in cancer. This xCT dependence may itself be a liability: system xc⁻ also imports selenocystine, and rather than protecting cells, high xCT sensitizes them to selenocystine-induced oxidative cytotoxicity (Tan et al., *Biochem J* 2023), nominating the xCT-high malignant pole as a candidate for selective engagement by selenocystine, a hypothesis we flag for future testing rather than examine here.

These opposing poles are not merely two routes to cystine but two spatially and mechanistically distinct systems, which is why xCT does not substitute for b^0,+^ despite an overlapping substrate range. b^0,+^ is an apical, brush-border amino-acid exchanger, the rBAT/b0,+AT system whose loss causes cystinuria through failed luminal cystine reabsorption in kidney and small intestine (Fotiadis et al., *Mol Aspects Med* 2013). It is therefore positioned to capture cystine and selenocystine directly from the gut lumen, i.e., from the diet. System xc⁻ is instead a cystine–glutamate antiporter that draws on the extracellular, non-luminal cystine pool, with a substantial non-canonical role at the lysosomal membrane (Zhou et al., *Cell* 2025); it is not a brush-border uptake system. Consistent with this, xCT is expressed at very low levels in healthy enterocytes (He et al., *bioRxiv* 2026), and its disease-associated induction arises precisely in cells that have lost normal absorptive identity, the dedifferentiated LND state in Crohn’s disease and the malignant colonocyte, where apical–basolateral polarity is already compromised. The thiol pole is thus not a re-routing of dietary selenium uptake but a stress/NRF2-driven, non-luminal cystine program layered on a depolarized epithelium, and b^0,+^ remains the functionally unique, diet-facing selenium-supply node that xCT cannot replace regardless of its capacity to carry selenocystine as a cystine analog.

Several boundaries apply. The disease-associated epithelium of the integrated atlas derives from a small number of donors and source studies, which is why the cell-state mechanism is established in the independent, severity-graded cohort; the two cohorts are complementary in breadth and depth. The ferroptosis state is inferred transcriptionally, and the inflammation-to-selenium-pole link is supported by donor-paired perturbation data in small-intestinal enteroids, IFNγ (Crohn’s-disease cohort significant, non-IBD cohort concordant but underpowered) and TNF (two paired datasets, selenoprotein-arm suppression with the b^0,+^ transporter spared), though this evidence, like IL-17’s line-variable signal, derives largely from a single laboratory (Methods) and is bulk rather than single-cell; none of it is yet backed by direct functional measurement in an isogenic system. A direct functional test is therefore the critical next step (Future directions, below). The transcriptional, proteomic and network-based evidence in this manuscript should accordingly be read as convergent and motivating, rather than yet confirming, the proposed causal supply-to-defense link; in particular, these data cannot exclude that the losses of *GPX4* and of b^0,+^ are parallel consequences of the same dedifferentiation program rather than causally linked, nor that *GPX4* falls through a cell-intrinsic, oxidant-driven translational block independent of substrate supply, alternatives the isogenic functional test below is designed to discriminate. The treatment-response correlate is likewise exploratory. Finally, this is a transcriptomic study and does not resolve protein subcellular localization; the apical/luminal-versus-non-luminal and lysosomal distinctions drawn between b^0,+^ and xCT above are interpretations grounded in prior literature rather than localization measurements made here.

### Future directions

The central prediction, that loss of b^0,+^ is directly sufficient to trigger the selenium→thiol switch and ferroptosis priming, is testable in an isogenic system. We outline a CRISPR *SLC7A9* knockout in polarized Caco-2 (which differentiate to a small-intestinal absorptive phenotype expressing b^0,+^; Fig 6B) versus an isogenic wild-type control, read out by (i) the selenoprotein/selenocysteine-incorporation module and the NRF2/xCT program (RNA-seq), (ii) the *GPX4*/*SELENOP*/xCT/driver proteome, cross-referenced to PXD036591, and (iii) ferroptosis sensitivity (BODIPY-C11 lipid peroxidation; RSL3/erastin dose–response). Because standard media supply selenium as selenite, which enters b^0,+^-independently, the decisive comparison uses selenocystine as the limiting selenium source on a selenium-depleted background, with a selenite arm as a b^0,+^-independent rescue control; a lighter cytokine-challenge arm (IFNγ/TNF/IL-17 on wild-type monolayers) would test whether inflammatory stimulation phenocopies genetic b^0,+^ loss. This design discriminates the alternatives above: a knockout that selectively collapses *GPX4* under selenocystine yet is rescued by selenite (a b^0,+^-independent selenium source) would establish a substrate-supply limitation, whereas persistent *GPX4* loss under selenite would implicate a *GPX4*-intrinsic translational block; and comparing the knockout with cytokine challenge in otherwise-differentiated monolayers separates supply-node loss from a dedifferentiation-parallel change. For these contrasts to be interpretable, knockout and wild-type lines must reach equivalent polarization and enterocyte identity (transepithelial resistance, tight-junction and identity markers), so that any *GPX4* difference is not itself secondary to impaired differentiation.

Clinically, the model frames a selenium–ferroptosis axis on which inflammatory and malignant intestinal disease occupy opposite positions. The actionable line from this work runs through the inflammatory pole: because the loss is substrate-supply–linked and graded by severity, restoring the selenium pole, via selenium/selenocystine availability, b^0,+^ support, or upstream cytokine control, is a candidate route to limiting the ferroptosis-prone dedifferentiated state in Crohn’s disease, contingent on the supply-node causality that the functional test above is designed to establish. The same axis may also carry prognostic information: patients whose epithelium had already switched toward the thiol pole at baseline responded less well to subsequent anti-TNF therapy (Fig 7C), so the supply-node state that grades disease severity may also track treatment response, an exploratory association, drawn from small bulk cohorts, that would require prospective validation before any clinical use. Translating this into nutritional, genetic and preventive strategies for Crohn’s disease is the subject of ongoing work.

## Methods

### Datasets

#### Pan-GI Atlas Extended

(Oliver et al., *Nature* 2024; https://gutcellatlas.org) provided the primary cross-disease atlas: 72,356 small-intestinal and 96,675 large-intestinal epithelial cells spanning healthy, Crohn’s disease, paediatric IBD, ulcerative colitis and colorectal cancer donors, integrated from 25 source studies with automated quality control (scAutoQC). Disease-group, tissue-microenvironment (sample_category), organ/segment and fine-grained cell-type annotations were taken from the atlas metadata; gene symbols were resolved from Ensembl IDs via the atlas feature_name/gene_symbols field.

#### GSE266546

(Vanderbilt Gut Cell Atlas; Li et al., *Nat Commun* 2024) provided an independent, severity-graded Crohn’s disease cohort: inDrops single-cell RNA-seq of terminal ileum and ascending colon from 65 Crohn’s disease and 18 non-IBD donors, with histological grading (Control; inactive: Normal_CD/Quiescent; active: Mild/Moderate/Severe) and author cell-type annotations including the LND (large non-differentiated) epithelial state. Raw counts and annotation files were obtained from GEO and rebuilt into a sparse AnnData object; counts were total-count normalized (10⁴) and log1p-transformed.

#### GSE16879

(Arijs et al., *PLoS One* 2009) provided an independent bulk-tissue validation cohort: Affymetrix HG-U133 Plus 2.0 profiling of mucosal biopsies from Crohn’s disease (ileal and colonic), ulcerative colitis (colonic) and control tissue, before and 4–6 weeks after first infliximab infusion, with endoscopic/histologic response classification. Probe intensities were collapsed to genes (maximum probe per gene) and log₂-transformed.

#### PXD036591

(Alfredsson et al., *Proteomics* 2023) provided an independent public proteome: isobaric-labeling (TMT) quantitative proteomics of FACS-purified EpCAM⁺ epithelial and immune cell fractions from Crohn’s disease and control intestinal biopsies. The EpCAM⁺ epithelial channels (3 Crohn’s-disease, 4 control) were re-analyzed for the b^0,+^/selenoprotein/ferroptosis/NRF2 protein panel reported in Results (Fig 6A); gene-symbol fields (frequently multi-name, semicolon- or slash-delimited) were split and alias-matched, and log₂ fold-change and Welch’s t-test were computed per protein on mean channel intensities (Crohn’s-disease versus control), both pooled and per-patient.

#### GSE162633

(Caco-2 differentiation RNA-seq) provided an independent public validation of the Caco-2 cellular model (Fig 6B): bulk transcriptomes of Caco-2 monolayers at day 8 (immature) versus day 16 (polarized), *n* = 4 per timepoint. Because this is a two-timepoint, four-replicate design rather than a maturation trajectory, analysis was restricted to effect direction and magnitude: per-gene log₂ fold-change of day-16-versus-day-8 group means, exact two-sided Mann–Whitney *U*, and rank-biserial effect size; no partial-correlation or maturation-independence claim was made (the exact Mann– Whitney floor at *n* = 4 per group is *P* = 0.029). The b^0,+^ (*SLC3A1/SLC7A9*), selenoprotein-program (*SELENOP/SEPHS2/GPX4/MSRB1*) and xCT (*SLC7A11*) panels were scored to confirm co-induction of the selenium pole with the thiol pole quiescent as the monolayer polarizes (Fig 6B).

#### Cytokine-perturbation screen

Public GEO transcriptomic datasets of human intestinal epithelium under cytokine or disease perturbation were retrieved by a documented search (GEO DataSets; a general intestinal-epithelium query returning 86 human RNA-seq Series and a small-intestine–restricted query returning 35; full Boolean queries and retrieval date in the archived search_strategy.md). Two sample-level inclusion criteria were applied, both critical for the b^0,+^/selenoprotein question: (i) a **small-intestinal** epithelial model, b^0,+^ (*SLC3A1*/*SLC7A9*) is a small-intestinal brush-border transporter, near-absent in colon, so colonic models (colonoids, T84) cannot test its regulation and were used only to assess the tissue-general xCT response, and (ii) **chronic** cytokine exposure (≥ 24 h; short 3-h challenges excluded). The datasets meeting both and directly interrogating the b^0,+^/selenoprotein program were small-intestinal enteroid cytokine experiments from one group (Hong and colleagues, Samsung Medical Center): IFNγ (GSE290417, ten Crohn’s-disease-derived lines; GSE290418, six non-IBD lines, one study, Lee et al., *Front Immunol* 2025) and TNF (GSE174792, n = 16 donor pairs; GSE153715, n = 12 pairs, experimentally and temporally distinct studies). These are independent sample sets but share a single laboratory, a limitation stated in Results. Per-gene log₂ fold-changes were computed from NCBI-recount TPM or submitter count matrices (Entrez-GeneID–indexed to avoid symbol-mapping error), CPM-normalized, and tested by donor-paired Wilcoxon signed-rank with Benjamini–Hochberg correction across the b^0,+^/selenoprotein panel. IL-17 (GSE182453, unpublished small-intestinal deposit) gave a line-variable signal, interpretable only in the one line with a robust b^0,+^ baseline, and is reported as consistent only. Datasets failing the criteria were excluded with documented reason: colonic tissue (e.g. GSE190634, GSE190705, GSE243230, GSE311760, GSE282444: b^0,+^ not expressed; xCT-only readout), short exposure (GSE146893, 3 h), a differentiation-medium rather than cytokine contrast (GSE202777, catalogued as ‘TNF’ but comprising only a WRN-concentration gradient), single-cell modality (GSE153802, GSE189423), or different species (GSE282271, mouse).

Independence of GSE266546 from the Pan-GI atlas was confirmed by the absence of shared donors, samples, source studies and sequencing chemistry (inDrops versus 10x throughout).

### Statistical framework

All disease-versus-control comparisons were performed on per-donor (equivalently per-sample, since each subject contributed a single sample passing the cell-count threshold) pseudobulk values, detection rate and/or mean expression among the relevant absorptive population, compared by two-sided Mann–Whitney U with Benjamini–Hochberg correction across readouts within an analysis. Cell-state (LND-versus-mature) contrasts used the donor-paired log₂ fold-change of mean normalized expression in LND versus the Mature Enterocyte cluster, with Wilcoxon signed-rank significance; this donor-level metric was applied uniformly to all LND-versus-mature estimates reported. Spearman correlation was used for severity trends, for gene–gene relationships (e.g., *SLC7A9*– *SELENOP*), and for relating epithelial readouts to immune-cell abundance or cytokine signatures. b^0,+^ co-expression was defined as cells positive for both *SLC7A9* and *SLC3A1* (or, as a continuous readout, the mean log-normalized expression of the two genes); the xCT complex was defined analogously from *SLC7A11/SLC3A2*. Maturation was scored two ways, for two distinct purposes. For the per-donor maturation comparison and the tercile stratification of the b^0,+^ deficit (Fig 2A,B), a scanpy score_genes score over canonical villus-enterocyte/absorptive markers (*APOA1*, *APOA4*, *APOB*, *FABP1/2/6*, *ALPI*, SI, *ANPEP*, *RBP2*, *SLC2A2*, *TREH*, *APOC3*, *MTTP*) was used. For the *SLC7A9*–*SELENOP* coupling (Fig 2C) we reproduced the method of He et al. (*bioRxiv* 2026), a 10-marker rank-normalized maturation score (*ALPI*, *FABP1*, *APOA1*, *APOA4*, SI, *CA1*, *CA2*, *CEACAM5*, *CEACAM6*, *KRT20*) with intestinal segment as covariates in a partial Spearman correlation across enterocytes, so that the disease result is directly comparable to the healthy-tissue coupling reported there; correlations were computed within each disease group and are accordingly lower in magnitude than the pooled cross-cohort value reported in the companion study. Selenoprotein, ferroptosis, NRF2-target and IFNγ-response modules were scored as the mean of z-scored member-gene expression, with genes undetected in a given dataset omitted from that dataset’s score. Proteomic comparisons (PXD036591) used Welch’s t-test on channel-level intensities given the small, unequal group sizes (n = 3 vs 4) and are reported as exploratory, without multiple-testing correction, alongside per-patient values to make between-patient heterogeneity explicit.

### Pan-GI Atlas Extended: donor-level reanalysis

Because cell-level significance testing in a multi-donor, multi-study atlas is subject to pseudoreplication, treating non-independent cells within a donor as independent replicates inflates significance and biases toward highly expressed genes (Squair et al., *Nat Commun* 2021), all Pan-GI atlas comparisons were re-derived at the donor level rather than the cell level. Disease epithelium was first audited for donor count, cell count per donor and source study of origin. The core Crohn’s-disease-versus-healthy b^0,+^ contrast used all Crohn’s-donor small-intestinal enterocytes (control_vs_disease = crohns_disease) versus organ-donor/control enterocytes, restricted to donors with ≥20 absorptive cells, with detection rate and continuous mean b^0,+^ score compared by donor-level Mann–Whitney U; three sensitivity checks were run in parallel: cell-count thresholds of 10/20/30 per donor, continuous versus detection-rate readouts, and a strict actively-inflamed-only definition versus all Crohn’s-disease donors, to confirm the result was not an artifact of any single analytic choice. Paediatric IBD was contrasted against age-matched non-IBD controls using the same donor-level framework. To test whether the b^0,+^ deficit reflected passive dedifferentiation, donor-level maturation scores were compared between disease and control enterocytes, and the b^0,+^ contrast was repeated within maturation terciles (and, as a feasibility check, deciles, found to be under-powered at the donor level and not used for inference). The *SLC7A9*–*SELENOP* relationship was assessed across enterocytes by partial Spearman correlation adjusting for maturation and intestinal segment (reproducing the companion study’s method; see Statistical framework), within both control and Crohn’s-disease groups, alongside the raw pseudobulk correlation.

To connect the LND cell state (identified and characterized directly in GSE266546, which lacks a native LND annotation for the Pan-GI atlas) to the predominantly healthy Pan-GI atlas, a maturation/inflammation transfer signature was built from the top genes up-regulated in LND versus mature enterocytes in GSE266546, explicitly excluding b^0,+^, selenoprotein and xCT pathway genes to avoid circular reasoning. This signature was scored in Pan-GI atlas small-intestinal enterocytes to define an LND-high/LND-low split, which was then tested for (i) expansion in inflamed versus non-pathological sample regions (using sample_category rather than diagnostic label, since paediatric IBD enterocytes did not show b^0,+^ suppression and diagnosis alone would conflate inflammatory state with disease label), (ii) lower b^0,+^ and *GPX4* expression than LND-low cells, and (iii) persistence of b^0,+^ suppression within maturation deciles.

### GSE266546: severity-graded cohort analysis

Disease and cell-state effects were assessed on absorptive (enterocyte/colonocyte) cells aggregated to per-sample pseudobulk means (62 terminal-ileum samples from 62 patients passing the cell-count threshold; a patient-random-intercept mixed model was therefore not required for the terminal-ileum severity analyses, and was nonetheless run as a robustness check, see below). LND-versus-mature comparisons contrasted LND with the Mature Enterocyte cluster within active disease using the donor-paired log₂ fold-change described above. Per-sample Spearman correlations related the epithelial selenium program to the abundance of inflammatory immune cells (neutrophils and recruited macrophages, as a fraction of all cells) and to an epithelial IFNγ-response signature (scored with scanpy score_genes). The selenocysteine-incorporation machinery (*SEPHS1/2*, *SECISBP2*, *EEFSEC*, *PSTK*, *SEPSECS*) and the two b^0,+^ subunits were compared between active and control samples and between LND and mature enterocytes (Mann–Whitney U). Cytokine producers were mapped by per-cell-type mean expression and detection rate of IFNG, IL17A/F, TNF, IL1B, IL22, IL6 and OSM across immune cell types; the per-sample immune-compartment cytokine signal and producer-cell fractions (T cells, macrophages, neutrophils) were correlated with the epithelial switch readouts. The full selenoproteome (23 of 25 members detected) was profiled along the same two axes: per-sample pseudobulk Spearman against severity, and LND-versus-mature log₂ fold-change, with each member annotated by the canonical selenium-deficiency hierarchy (high-versus low-priority) to distinguish selective from global loss. Enterocyte-identity transcription factors (*HNF4A*, *HNF4G*, *CDX2*, *GATA4*, *KLF5*) and an NRF2-target program (*NQO1*, *GCLC*, *GCLM*, *SLC7A11*, *TXNRD1*, *GSR*, *ME1*, *G6PD*, *GPX2*, used as a transcriptional proxy for NRF2 activity, since *NFE2L2* transcript itself is a poor readout of its post-translationally regulated activity) were related to the b^0,+^ score, *GPX4* and xCT by per-sample Spearman correlation and tested for LND-versus-mature change. Candidate selenium-uptake receptors (*LRP1/2/5/6/8*, *CUBN*, *AMN*) were profiled along the same severity and LND-versus-mature axes and by per-sample detection rate, to compare directly with detection-rate findings in the Pan-GI atlas; co-expression with the b^0,+^ score was additionally assessed within mature enterocytes by depth-adjusted partial Spearman correlation. Early-versus-late LND subpopulations were scored and related to the redox-switch readouts and to histological severity. Lineage-specific robustness was assessed by repeating severity correlations separately within each annotated cell type (stem/transit-amplifying, absorptive, LND, goblet, immune, stromal), and by a patient-level mixed-effects model with a random intercept as an additional check on the pseudobulk severity trends. Clinical metadata (CDAI and medication exposure) from the original study’s supplementary table were merged by sample, and robustness to treatment was assessed by repeating the severity and infiltrate correlations within anti-TNF–naive and biologic-naive subsets. Cell-cycle phase was scored with scanpy score_genes_cell_cycle using the standard Tirosh S- and G2/M-marker lists; a cell was called cycling if assigned to S or G2M, with stem/transit-amplifying cells as an internal positive control, and the LND-versus-mature cycling fraction was compared donor-paired within active disease (Wilcoxon signed-rank) and re-checked in sequencing-depth-matched cells (total counts restricted to the pooled interquartile band). Coordinated identity loss was tested by self-contained prerank GSEA (weighted enrichment score with a gene-set permutation null and Benjamini–Hochberg FDR) of a mature-enterocyte identity gene set against two independent rankings: the donor-paired LND-versus-mature log₂FC and the per-sample severity Spearman metric. The absorptive-lineage trajectory was built on the 2,000 most variable genes with Harmony batch integration over sample of origin (harmonypy 2.0; Korsunsky et al., *Nat Methods* 2019), followed by PAGA connectivity and diffusion pseudotime (root = the lowest-maturation stem cell); the LND topology was confirmed stable across uncorrected, ComBat and Harmony batch treatments. The severity-graded redox switch (Fig 3B) was quantified as a per-sample pseudobulk composite score (mean z-scored thiol-arm minus mean z-scored selenium-arm expression) rather than by set-level enrichment testing: at pseudobulk resolution the individual selenium- and thiol-pole gene sets are too sparse (k = 3–5 detected members after per-sample filtering, with *SLC7A11* frequently below the detection floor) for a stable enrichment statistic, whereas the composite aggregates the same members into a full-power donor-level readout, so set-level GSEA of these small pole sets is under-powered by construction and is not used to test the switch.

### GSE16879: bulk validation and anti-TNF response

To validate the switch on an independent platform, active (pre-treatment) Crohn’s-disease ileal and colonic and ulcerative-colitis colonic samples were each compared with segment-matched controls (Mann–Whitney U per gene), and a composite switch-score was defined per sample as the mean z-scored expression of the induced arm (*SLC7A11*, *GPX2*, *GCLC*, *AIFM2*) minus the homeostatic arm (*SLC7A9*, *GPX4*, *SELENOP*). For response prediction, baseline (pre-infliximab) Crohn’s-disease ileal samples from subsequent responders (n = 8) and non-responders (n = 10) were compared by Mann– Whitney U on two pre-specified switch readouts (the b^0,+^ score and *SLC7A11*/xCT), with rank-biserial effect sizes reported; patient-matched before/after changes were assessed by Wilcoxon signed-rank; and an exploratory ROC/AUC was computed for the composite switch-score, treated as hypothesis-generating given the small group sizes and nominal (uncorrected) *P* values.

### Cytokine-perturbation screen: replicate-structure audit and statistical framework

An initial automated screen classified per-gene direction (up/down/flat at |log₂FC| ≥ 0.25 of group means) across the retrieved datasets without regard to tissue, replicate structure or formal significance testing. On reaudit this was found to conflate independent biological replication with technical/passage replication of a single line for several datasets, which would overstate the evidential weight of the screen; the analysis reported here replaces that screen with an explicit replicate-structure audit and donor-appropriate statistics.

For each dataset, per-sample metadata (GEO characteristics fields) were used to determine the true unit of biological replication. Two datasets (GSE290417, GSE290418) had unambiguous multi-donor paired designs (the same line profiled with and without IFNγ) and were re-analyzed with the donor-paired framework used throughout this study: per-donor log₂ fold-change (IFNγ versus each line’s matched untreated control) was computed for each core b^0,+^/selenoprotein gene and tested by two-sided Wilcoxon signed-rank across donors, with Benjamini–Hochberg correction applied across the genes tested within each dataset. A between-cohort comparison (Mann–Whitney U on the two datasets’ per-donor log₂ fold-changes) tested whether the Crohn’s-disease-derived cohort (GSE290417) showed a stronger IFNγ response than the non-IBD cohort (GSE290418) for each gene; none reached significance (P = 0.26–0.96), so no disease-specific IFNγ-hypersensitivity claim is made.

TNF was tested directly in two paired small-intestinal enteroid datasets from the same group (GSE174792, eight control + eight Crohn’s-disease donor pairs; GSE153715, jejunal and ileal lines ± TNF): per-donor log₂ fold-changes (TNF versus each line’s matched untreated control) were computed from CPM-normalized counts and tested by donor-paired Wilcoxon signed-rank across the b^0,+^/selenoprotein panel. GSE282444, initially catalogued as a TNF dataset, was reclassified as colonic (proximal/distal-colon organoids) and is therefore uninformative for b^0,+^, which is not expressed in colon; it is not used for the b^0,+^/selenoprotein claim. IL-17 (GSE182453) is small-intestinal but line-variable, interpretable only in the one line with a robust b^0,+^ baseline (a near-zero baseline destabilizes log₂FC, as also seen for *SLC7A9* generally), and is reported descriptively. GSE282271 (mouse IFNγ) was excluded (single pooled line; different species). Datasets failing the small-intestinal or chronic-exposure criteria, colonic models, short (3-h) challenges, differentiation-medium gradients, and single-cell deposits, were reviewed as context only; none produced concordant b^0,+^/SE-core downregulation, consistent with the co-suppression being a feature of cytokine signalling in small-intestinal epithelium rather than a generic epithelial-state change.

### Network-based virtual knockout

Network-based virtual knockout (scTenifoldKnk; Osorio et al., *Patterns* 2022) was performed on a gene-regulatory network constructed from normal (non-diseased) small-intestinal enterocytes, deliberately using physiological rather than diseased tissue so that the inferred regulatory structure would not be confounded by the disease-associated b^0,+^- to-xCT program switch. Each b^0,+^ subunit (*SLC3A1*, *SLC7A9*) was virtually deleted from the constructed network in turn; manifold alignment between the perturbed and wild-type networks identified differentially regulated genes, which were tested for gene-set enrichment (GSEA) against the selenoprotein gene set and, as an internal positive/sensitivity control, the ferroptosis gene set.

### Software

Analyses used Python 3.10 with scanpy, anndata, pandas, numpy, scipy (Mann–Whitney U, Spearman, Wilcoxon signed-rank, Welch’s t-test), statsmodels (Benjamini–Hochberg correction), matplotlib/seaborn (figures), harmonypy 2.0 (Harmony batch integration) and scTenifoldKnk (virtual knockout). A lightweight reproducibility package (paperb_repro) stages inputs, patches machine-specific paths, executes the numbered analysis scripts in a documented order, and audits expected outputs by checksum; the canonical LND-versus-mature effect sizes reported throughout come from this package’s lnd_vs_mature_canonical stage.

---

**Fig S1.**
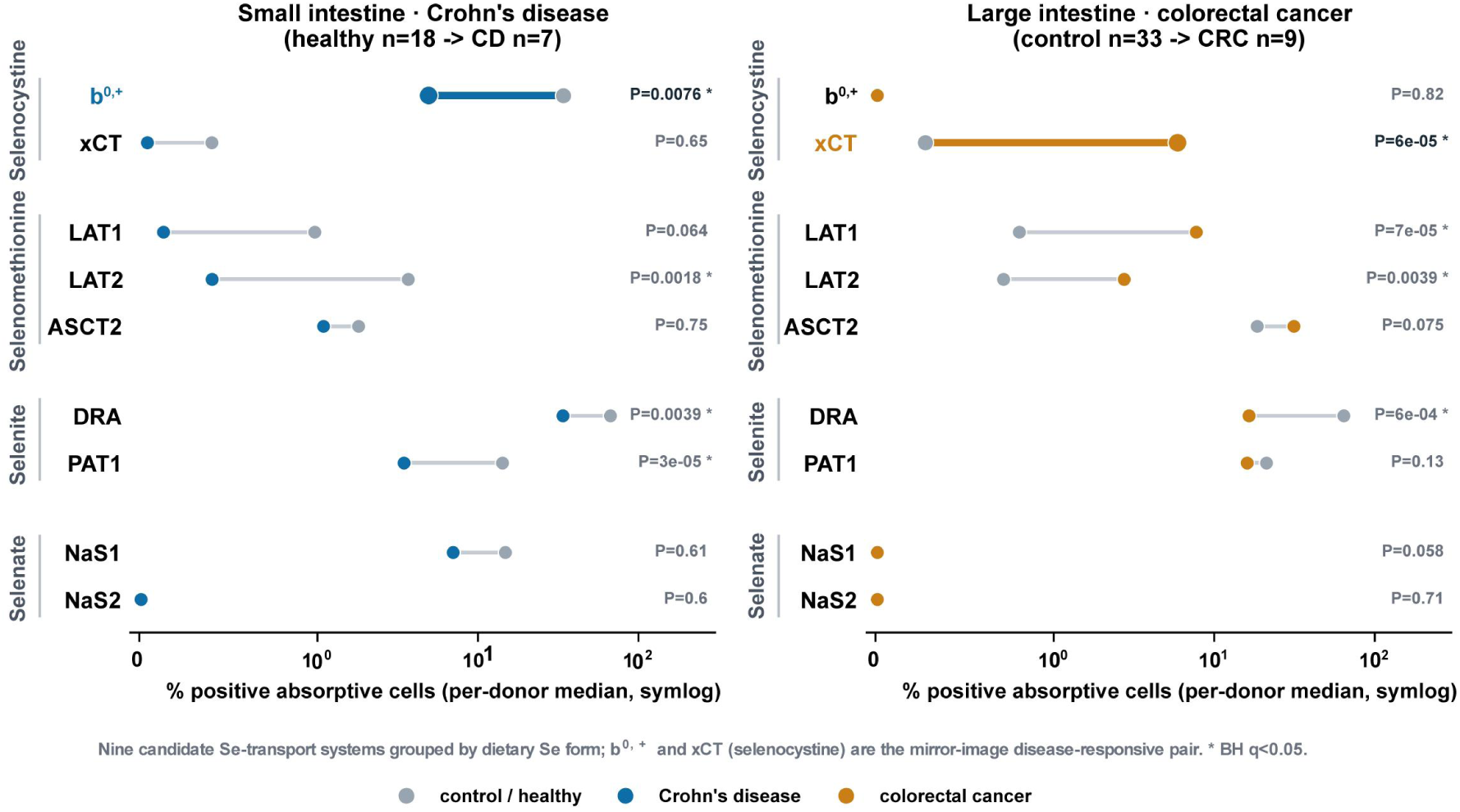
Donor-level disease contrast of all nine candidate selenium-transport systems across the four dietary selenium forms. Per-donor median % of absorptive cells positive for each system (heterodimers scored as co-detection of both subunits), healthy→adult CD in small intestine (n = 18 → 7; donor_disease grouping) and control→CRC in large intestine (n = 33 → 9); symlog axis; two-sided Mann–Whitney, Benjamini–Hochberg corrected (* q < 0.05). Among the two selenocystine systems, b^0,+^ is suppressed in Crohn’s small intestine and xCT induced in colorectal cancer, the mirror-image poles of the axis; transporters for the other dietary forms (selenomethionine: LAT1/LAT2/ASCT2; selenite: DRA/PAT1; selenate: NaS1/NaS2) change along distinct differentiation- and proliferation-associated programs (e.g. LAT1 co-induced with xCT in CRC; DRA/PAT1 lost with dedifferentiation). Cohort and positivity definitions match Fig 1D/E.

**Fig S2.**
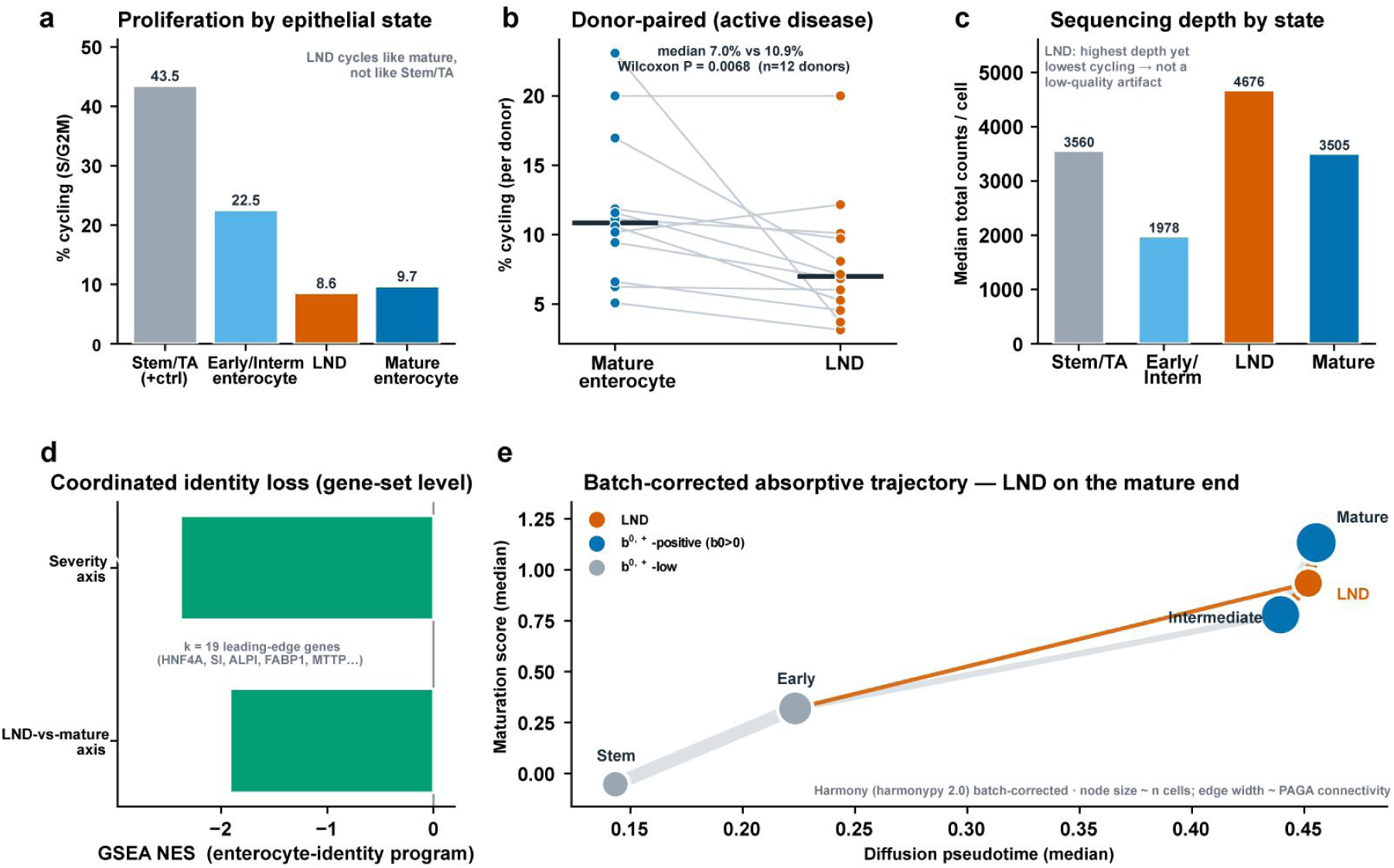
LND is a non-proliferative, gene-set-level dedifferentiated state on the differentiated end of the absorptive trajectory. Severity-graded Crohn’s cohort (GSE266546, terminal ileum); absorptive lineage plus stem/transit-amplifying cells. **(A)** Cell-cycle (S/G2M) fraction by epithelial state: LND (8.6%) cycles like mature enterocytes (9.7%), far below the stem/transit-amplifying positive control (43.5%). **(B)** Donor-paired cycling fraction within active disease (n = 12 donors): LND cycles less than mature enterocytes (median 7.0% vs 10.9%, Wilcoxon *P* = 0.007). **(C)** Per-state sequencing depth: LND carries the highest median total counts of any state, so its low cycling is not a low-depth/low-quality artifact. **(D)** Prerank GSEA of the mature-enterocyte identity program: coordinately down-regulated along both the LND-versus-mature axis (NES = −1.92, FDR = 0.009) and the independent histological-severity axis (NES = −2.38, FDR = 0.013; k = 19 leading-edge genes). **(E)** Harmony batch-corrected PAGA / diffusion-pseudotime trajectory of the absorptive lineage: nodes positioned by median pseudotime (x) and maturation score (y), sized by cell number, edges weighted by PAGA connectivity; LND sits at the mature end, most strongly connected to intermediate and mature enterocytes (0.63 / 0.79) and least to stem cells (0.09), a dedifferentiated offshoot of maturing enterocytes, not a stem-like state; topology consistent across uncorrected, ComBat and Harmony batch treatments.

**Fig S3.**
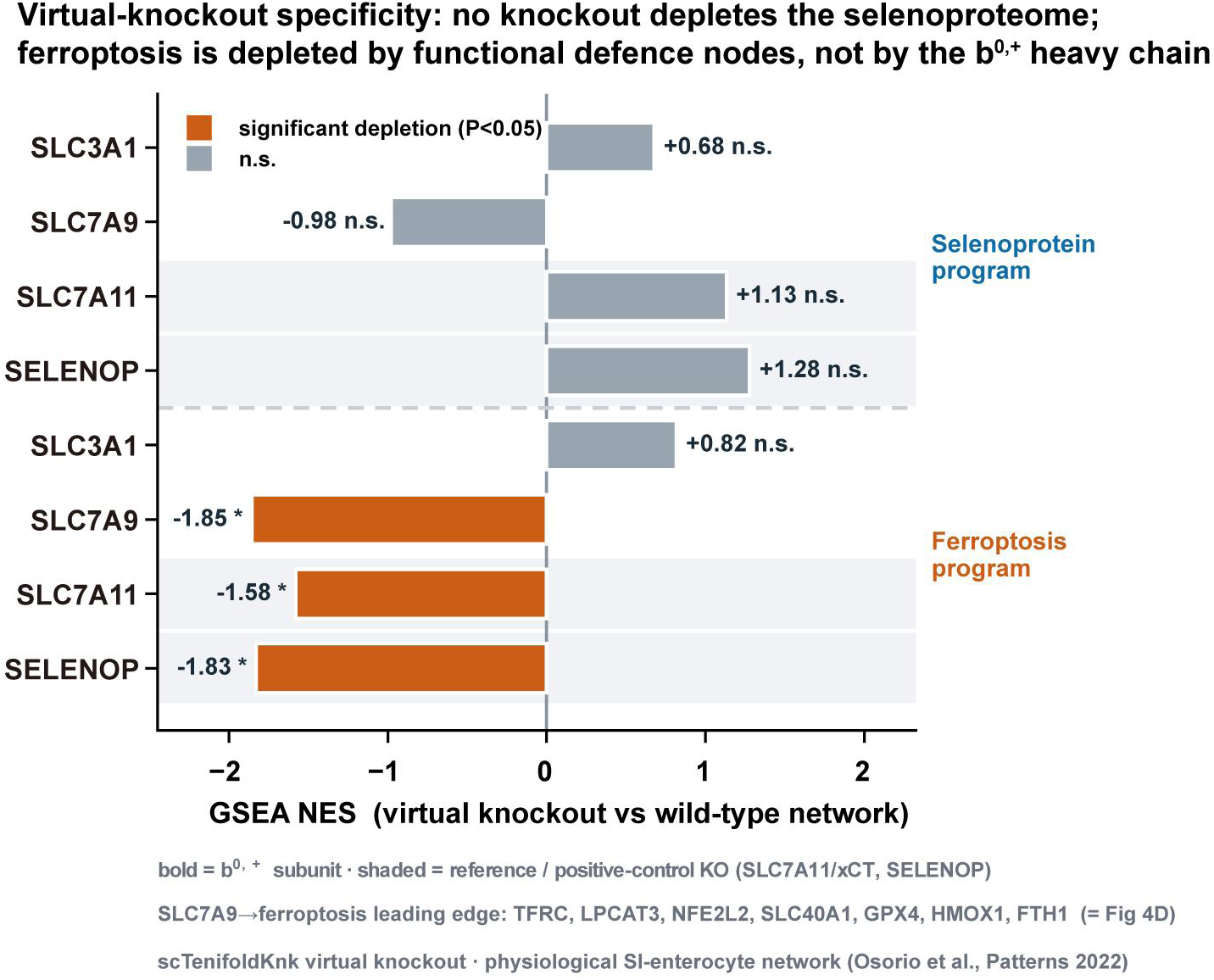
Virtual-knockout specificity: no knockout depletes the selenoproteome; the ferroptosis program is depleted by functional ferroptosis-defence nodes but not by the b^0,+^ heavy chain. scTenifoldKnk virtual knockout in a physiological small-intestinal enterocyte network, with GSEA of the perturbed-versus-wild-type network against the selenoprotein and ferroptosis gene sets, for the two b^0,+^ subunits (*SLC3A1* heavy chain, *SLC7A9* catalytic) and two reference/positive-control knockouts (*SLC7A11*/xCT and *SELENOP*, established ferroptosis-defence genes). None of the four knockouts depleted the selenoprotein set (all *P* > 0.16; *P* ≥ 0.49 for the two b^0,+^ subunits), arguing against a direct transcriptional b^0,+^→selenoprotein axis, even *SELENOP* knockout did not deplete the selenoprotein module in this transcriptional network. The ferroptosis set was depleted by the functional defence nodes *SLC7A9* (NES = −1.85, *P* = 0.017; leading edge including *GPX4*, *FTH1*, *SLC40A1*, *TFRC*), *SLC7A11* (NES = −1.58, *P* = 0.036) and *SELENOP* (NES = −1.83, *P* = 0.016), confirming that the network detects genuine ferroptosis-defence links, but not by the b^0,+^ heavy chain *SLC3A1* (NES +0.82, n.s.). The *SLC7A9* result is the supply→vulnerability link shown in Fig 4D. Bold = b^0,+^ subunit; shaded rows = reference KOs.

## Declarations

### Data availability

All datasets analysed in this study are publicly available; no new primary data were generated. Accessions and repositories:

- **Pan-GI Atlas Extended**, integrated cross-disease single-cell atlas (Oliver et al., *Nature* 2024); available via the Gut Cell Atlas (https://gutcellatlas.org).
- **GSE266546**, severity-graded Crohn’s-disease single-cell cohort (Li et al., *Nat Commun* 2024); NCBI GEO.
- **GSE16879**, bulk mucosal transcriptomes with infliximab-response annotation (Arijs et al., *PLoS One* 2009); NCBI GEO.
- **PXD036591**, FACS-purified EpCAM⁺ epithelial TMT proteome (Alfredsson et al., *Proteomics* 2023); PRIDE/ProteomeXchange.
- **GSE162633**, Caco-2 polarization (day 8 vs day 16) RNA-seq; NCBI GEO.
- **GSE290417 / GSE290418**, small-intestinal enteroid IFNγ RNA-seq (Lee et al., *Front Immunol* 2025); NCBI GEO.
- **GSE174792 / GSE153715**, small-intestinal enteroid TNF RNA-seq; NCBI GEO.
- **GSE182453**, small-intestinal enteroid IL-17 RNA-seq; NCBI GEO.

Processed, per-figure source tables generated in this study are available from the corresponding author on reasonable request, and will be released publicly together with the analysis code upon peer-reviewed publication.

### Code availability

Analysis code, the numbered per-Results verification scripts and notes, and a reproducibility package (paperb_repro) that stages inputs, patches machine-specific paths, and re-executes the analyses in a documented order with checksum-audited outputs are available from the corresponding author on reasonable request.

### Ethics approval and consent

This study is a secondary computational analysis of previously published, publicly available, de-identified human datasets and did not require additional ethical approval or consent; the original studies obtained their respective institutional approvals and participant consent.

### Declaration on the use of generative AI tools

During the preparation of this manuscript, the authors used AI to improve language and readability, as well as to assist with code checking and optimization. All outputs were critically reviewed and validated by the authors. The authors take full responsibility for the final content, including the correctness and integrity of the code and the overall manuscript.

### Author contributions (CRediT). Xiaobai He

Conceptualization, Methodology, Investigation, Formal analysis, Writing (original draft), Visualization, Funding acquisition. **Kundu Zhong:** Methodology, Investigation, Data curation, Formal analysis. **Yiyi Pu:** Methodology, Data curation. **Hui Hu:** Methodology, Investigation. **Yaoqiang Du:** Methodology, Data curation. **Xiaopan Chen:** Methodology, Writing (review& editing). **Linjie Chen:** Conceptualization, Formal analysis, Writing (review & editing), Supervision, Project administration.

## Acknowledgements

The authors thank the investigators who generated and publicly shared the datasets analysed here.

## References

Alfredsson J, Fabrik I, Gorreja F, et al. Isobaric labeling-based quantitative proteomics of FACS-purified immune cells and epithelial cells from the intestine of Crohn’s disease patients reveals proteome changes of potential importance in disease pathogenesis. Proteomics. 2023;23(5):e2200366. doi:10.1002/pmic.202200366 [PXD036591]

Arijs I, De Hertogh G, Lemaire K, et al. Mucosal gene expression of antimicrobial peptides in inflammatory bowel disease before and after first infliximab treatment. PLoS One. 2009;4(11):e7984. doi:10.1371/journal.pone.0007984 [GSE16879]

Dixon SJ, Lemberg KM, Lamprecht MR, et al. Ferroptosis: an iron-dependent form of nonapoptotic cell death. Cell. 2012;149(5):1060–1072. doi:10.1016/j.cell.2012.03.042

Fotiadis D, Kanai Y, Palacín M. The SLC3 and SLC7 families of amino acid transporters. Mol Aspects Med. 2013;34(2-3):139–158. doi:10.1016/j.mam.2013.06.008

GSE182453. RNA sequencing for enteroids treated with different concentrations of IL-17. Hong SN, Lee C (Samsung Medical Center). NCBI Gene Expression Omnibus, deposited 2021. No associated peer-reviewed publication at time of writing.

GSE282444. Epigenetic memory of intestinal epithelial cells in inflammatory bowel disease [RNA-seq]. Druliner BR, Hamdan F (Mayo Clinic). NCBI Gene Expression Omnibus, deposited 2024. No associated peer-reviewed publication at time of writing.

Hangauer MJ, Viswanathan VS, Ryan MJ, et al. Drug-tolerant persister cancer cells are vulnerable to GPX4 inhibition. Nature. 2017;551(7679):247–250. doi:10.1038/nature24297

He X, Zhong K, Yang W, Cao J, Chen X, Chen L. The b^0,+^ amino acid transporter defines a selenium-utilization enterocyte program in the human intestine. bioRxiv. 2026 May 30. doi:10.64898/2026.05.27.728150 [companion paper]

Huangfu S, Zheng J, He J, et al. Protective role of seleno-amino acid against IBD via ferroptosis inhibition in enteral nutrition therapy. iScience. 2024;27(9):110494. doi:10.1016/j.isci.2024.110494

Hughes DJ, Fedirko V, Jenab M, et al. Selenium status is associated with colorectal cancer risk in the European prospective investigation of cancer and nutrition cohort. Int J Cancer. 2015;136(5):1149–1161. doi:10.1002/ijc.29071

Ingold I, Berndt C, Schmitt S, et al. Selenium utilization by GPX4 is required to prevent hydroperoxide-induced ferroptosis. Cell. 2018;172(3):409–422.e21. doi:10.1016/j.cell.2017.11.048

Jiang X, Stockwell BR, Conrad M. Ferroptosis: mechanisms, biology and role in disease. Nat Rev Mol Cell Biol. 2021;22(4):266–282. doi:10.1038/s41580-020-00324-8

Koppula P, Zhuang L, Gan B. Cystine transporter SLC7A11/xCT in cancer: ferroptosis, nutrient dependency, and cancer therapy. Protein Cell. 2021;12(8):599–620. doi:10.1007/s13238-020-00789-5

Korsunsky I, Millard N, Fan J, et al. Fast, sensitive and accurate integration of single-cell data with Harmony. Nat Methods. 2019;16(12):1289–1296. doi:10.1038/s41592-019-0619-3

Lee C, Kim JE, Cha YE, et al. IFN-γ-induced intestinal epithelial cell-type-specific programmed cell death: PANoptosis and its modulation in Crohn’s disease. Front Immunol. 2025;16:1523984. doi:10.3389/fimmu.2025.1523984 [GSE290417, GSE290418]

Li J, Simmons AJ, Hawkins CV, et al. Identification and multimodal characterization of a specialized epithelial cell type associated with Crohn’s disease. Nat Commun. 2024;15(1):7204. doi:10.1038/s41467-024-51580-7 [GSE266546]

Nakajima-Koyama M, Kabata M, Lee J, et al. The balance between IFN-γ and ERK/MAPK signaling activities ensures lifelong maintenance of intestinal stem cells. Cell Rep. 2025;44(3):115286. doi:10.1016/j.celrep.2025.115286 [GSE282271]

Oliver AJ, Huang N, Bartolome-Casado R, et al. Single-cell integration reveals metaplasia in inflammatory gut diseases. Nature. 2024;635(8039):699–707. doi:10.1038/s41586-024-07571-1

Osorio D, Zhong Y, Li G, et al. scTenifoldKnk: an efficient virtual knockout tool for gene function predictions via single-cell gene regulatory network perturbation. Patterns (N Y). 2022;3(3):100434. doi:10.1016/j.patter.2022.100434

Rayman MP. Selenium and human health. Lancet. 2012;379(9822):1256–1268. doi:10.1016/S0140-6736(11)61452-9

Speckmann B, Steinbrenner H. Selenium and selenoproteins in inflammatory bowel diseases and experimental colitis. Inflamm Bowel Dis. 2014;20(6):1110–1119. doi:10.1097/MIB.0000000000000020

Squair JW, Gautier M, Kathe C, et al. Confronting false discoveries in single-cell differential expression. Nat Commun. 2021;12(1):5692. doi:10.1038/s41467-021-25960-2

Tan SLW, Tan HM, Israeli E, et al. Up-regulation of *SLC7A11*/xCT creates a vulnerability to selenocystine-induced cytotoxicity. Biochem J. 2023;480(24):2045–2058. doi:10.1042/BCJ20230317

Vinceti M, Filippini T, Del Giovane C, et al. Selenium for preventing cancer. Cochrane Database Syst Rev. 2018;1(1):CD005195. doi:10.1002/14651858.CD005195.pub4

Xu M, Tao J, Yang Y, et al. Ferroptosis involves in intestinal epithelial cell death in ulcerative colitis. Cell Death Dis. 2020;11(2):86. doi:10.1038/s41419-020-2299-1

Zhou N, Chen J, Hu M, et al. *SLC7A11* is an unconventional H+ transporter in lysosomes. Cell. 2025;188(13):3441–3458.e25. doi:10.1016/j.cell.2025.04.004

Zhang X, Ma Y, Ma J, Yang L, et al. Glutathione Peroxidase 4 as a Therapeutic Target for Anti-Colorectal Cancer Drug-Tolerant Persister Cells. Front Oncol. 2022;12:913669. doi:10.3389/fonc.2022.913669. PMID: 35719967.

